# Mathematical Models of Retinal Drug Delivery

**DOI:** 10.1101/2024.12.17.629033

**Authors:** Paul A. Roberts, Chloe N. Thomas, Gabriel Bellamy Plaice, James A. Roberts, Marie-Christine Jones, James W. Andrews, Lisa J. Hill

**Affiliations:** Centre for Systems Modelling and Quantitative Biomedicine, University of Birmingham, Birmingham, United Kingdom; Department of Optometry and Visual Sciences, City St George’s, University of London, London, United Kingdom; Department of Biomedical Sciences, University of Birmingham, Birmingham, United Kingdom; School of Pharmacy, University of Birmingham, Birmingham, United Kingdom; School of Mathematics, University of Birmingham, Birmingham, United Kingdom

**Author notes:** Corresponding author: E-mail address (Paul A. Roberts). **Grant information:** This study was funded through the Centre for Systems Modelling and Quantitative Biomedicine by the University of Birmingham Dynamic Investment Fund. P.A.R. was funded by the University of Birmingham Dynamic Investment Fund. C.N.T. was funded by Fight for Sight. G.B.P. was funded by the Kennedy Trust for Rheumatology Research. J.A.R. was funded by the Macular Society.

## Abstract

**Purpose:** Wet age-related macular degeneration (AMD) causes vision loss when vascular endothelial growth factor (VEGF) stimulates blood vessel growth into the light-sensitive retina. Anti-VEGF treatments such as ranibizumab are currently administered to treat wet AMD via intravitreal injections, which are unpleasant, expensive and risk complications. We explored the efficacy of topically administered ranibizumab, with cell penetrating peptides (CPPs).

**Methods:** *Ex vivo* pig eyes were divided into 3 groups and treated with 1. topical or 2. intravitreal ranibizumab and CPP, or 3. intravitreal ranibizumab. ELISAs measured ranibizumab and VEGF concentrations in the aqueous and vitreous at 20 min, 40 min, 1 hr and 3.5 hr (*n* = 3, per group). An ordinary differential equation model was formulated to describe the evolving concentrations of ranibizumab, VEGF and their compounds in the tear, aqueous and vitreal compartments.

**Results:** Experimental — Topical: aqueous ranibizumab levels increased significantly, coincident with a significant drop in aqueous VEGF. Vitreal ranibizumab increased significantly, while vitreal VEGF remained constant. Intravitreal (with and without CPP): vitreal ranibizumab reached high concentrations, coincident with a significant drop in vitreal VEGF. Mathematical — topical treatment may provide sustained, moderate suppression of vitreal VEGF levels, while intravitreal treatment provides strong suppression which lessens between treatments.

**Conclusions:** CPP allows topical ranibizumab to penetrate the cornea but reduces ranibizumab availability and efficacy in neutralising VEGF for intravitreal treatment. Combined intravitreal/topical treatment presents a promising approach. Treatment efficacy would be enhanced if ranibizumab’s rate of binding to VEGF or tear residence time could be increased.

## Introduction

Intravitreal injections, the standard mode of drug administration for wet age-related macular degeneration (AMD), are unpleasant, expensive and risk complications. In this paper, we take a combined experimental-modelling approach, exploring the potential for topical administration to replace or augment intravitreal treatment.

AMD is a degenerative retinal disease, which leads to irreversible loss of central vision. Affecting approximately 196 million people worldwide, it is a growing health and economic burden, with 288 million cases expected by 2040. ^1^ AMD progresses through early, intermediate and late stages. ^2^ The late stage comes in two forms: dry and wet. ^2^ Dry AMD involves the degeneration of the neural retina, retinal pigment epithelium and choroid in a process known as geographic atrophy, while wet AMD is marked by choroidal neovascularisation (CNV), wherein, encouraged by vascular endothelial growth factor (VEGF), blood vessels from the choroidal vasculature undergo aberrant growth, penetrating and damaging the retina. ^2–5^

Wet AMD is currently treated with regular, invasive, intravitreal injections of anti-VEGF drugs, such as ranibizumab, bevacizumab and aflibercept (all biologicals), ^6^ which bind to and neutralise VEGF, ^6^ reducing CNV, and slowing vision loss. These injections are expensive, due, in part, to the need for regular medical appointments, and are associated with complications, including discomfort, endophthalmitis, retinal detachments and subconjunctival haemorrhages, resulting in suboptimal drug adherence from patients. ^7^ As such, there is an urgent clinical need to develop less invasive drug-delivery approaches. ^8^

Topical delivery provides a promising, non-invasive alternative, in the form of eye drops or drug-eluting contact lenses. ^9,10^ While promising, these delivery methods present new challenges, including the difficulty of transporting anti-VEGF molecules across the cornea (mostly because current anti-VEGF molecules are all fairly large), and the need to achieve therapeutic doses at the back of the eye, overcoming various barriers (e.g. aqueous-vitreous and vitreo-retinal interfaces) and clearance mechanisms (e.g. tear and aqueous clearance and dilution). ^11^ The first of these challenges may be overcome using cell-penetrating peptides (CPPs); positively-charged short peptide chains which serve as chaperones for therapeutics, enhancing their delivery through tissues. ^12,13^ Pre-clinical studies have demonstrated that CPPs may aid drug delivery to the retina, ^14^ including the delivery of ranibizumab and bevacizumab ^9^ (and to other structures with notoriously obstructive barriers such as the skin ^15^ and cell plasma membranes ^16^).

In this study, we explore the potential of topically delivered ranibizumab and a poly-arginine based CPP for the treatment of wet AMD, taking both experimental and mathematical approaches. Experimentally, we use an *ex vivo* porcine model to measure ranibizumab and VEGF concentrations over time, in the aqueous and vitreous, following topical or intravitreal administration (extending de Cogan et al. ^9^ which considered only a single time-point in porcine eyes and did not measure aqueous or VEGF concentrations). Mathematically, we develop an ordinary differential equation model to describe and predict the evolving concentrations of ranibizumab, VEGF and their compounds in the tear, aqueous and vitreous. Our model is based on that of Hutton-Smith et al. ^17^, with a number of important differences; most significantly, we include a tear compartment and a modified aqueous-vitreous exchange term to allow for the passage of ranibizumab, VEGF and their compounds in both directions across the aqueous-vitreous interface (where before they could move only in the vitreous to aqueous direction). Fitting this model to our experimental data, we use it to extrapolate to the *in vivo* human eye, predicting treatment efficacy for a range of treatment regimens and scenarios. (See reviews of mathematical models of ocular drug delivery ^18–21^ and of the retina ^22^ for details of previous modelling work in this area.)

Our model has the potential to accelerate the development and implementation of new ocular treatment approaches by providing an efficient and cost-effective means of testing various regimens. Further, our model can be straight-forwardly extended to predict the dynamics of other ocular drugs, including anti-VEGF treatments such as bevacizumab and aflibercept, or new and upcoming treatments for dry AMD such as pegcetacoplan and avacincaptad pegol. ^23–25^

## Methods

### Experimental methods

#### CPP formulation

Lyophilised CPPs, ∼6 amino acids long, were purchased from Genscript (SC1208) and reconstituted in sterile, nuclease-free water.

#### Ranibizumab formulation

1 mg of humanised anti-VEGF monoclonal antibody fragments (Ranibizumab) was purchased already constituted in PBS from MedChem Express (CAT: HY-P9951A-1mg; New Jersey, USA) and stored at −80 degrees Celsius.

#### Zeta potential

Ranibizumab (1 mg ml^−1^) and CPP (20–100 mg ml^−1^) solutions were prepared in water. Complexes were produced by mixing the antibody and CPP at various concentration (ranibizumab:CPP – 1:1, 1:10, 1:20, 1:25, 1:50, 1:100, 10:10, 10:25 and 10:50) and volume (ranibizumab vol.:CPP vol. – 1:1 starting with 10 or 50 µl ranibizumab, and 1:2, 1:3 and 1:5 starting with 10 µl ranibizumab) ratios in a 96-well plate at room temperature. The zeta potential of the complexes was assessed by electrophoretic light scattering on a Nanosizer ZS (Malvern Panalytical, Malvern, Worcestershire, UK) after dilution in aqueous sodium chloride to adjust conductivity (10 µL of complexes with NaClaq 5–7 mM). All measurements were performed at room temperature.

For most complexes, bi- or tri-modal distributions were observed. When CPP concentration was low (< 50 mg mL^−1^), results were more variable, potentially due to low complex concentration and, therefore, increased noise. As the CPP concentration was increased to 50 or 100 mg ml^−1^, measurements appeared more robust, with improved quality. For the 1:100 mg ml^−1^ complexes, average zeta potential values were slightly positive (ca. 4–6 mV), with the presence of strongly positive complexes (>15 mV) noted within some of the samples. The change in zeta potential observed following CPP addition suggests successful formation of the complexes.

A concentration ratio of 1:100 mg ml^−1^ was selected for the preparation of the complexes for the *ex vivo* experiments, which corresponds to a large molar excess of CPPs compared to the monoclonal antibody fragment. The zeta potential of the complexes was measured 24 hr after preparation. Zeta potential appeared stable within this time frame with values ca. 3–6 mV. Two populations were detected with zeta potentials ca. ±15 mV.

#### Administering treatment to *ex vivo* porcine eyes

A total of 39 fresh, unscalded *ex vivo* porcine eyes were delivered in a cooled box <24 hr after death. On arrival, the eyes were placed in phosphate-buffered saline after removing excess tissue. Eye diameters (length and height/width) were recorded using a calliper to inform geometrical parameters in the mathematical model. Eyes were placed in 6-well plates, supported with foam inserts, and 45 µl of therapeutics was applied either as a topical drop or via intravitreal injection.

The experimental groups are listed in Table 1 (with 12 eyes for each of the 3 treatment groups and 3 eyes for controls). Eyes received either a topical eye drop of ranibizumab (1 mg ml^−1^) and CPP (100 mg ml^−1^); an intravitreal injection of ranibizumab (1 mg ml^−1^) and CPP (100 mg ml^−1^); or an intravitreal injection of ranibizumab alone (1 mg ml^−1^). The remaining three eyes received an intravitreal injection of sterile water as a control (see Fig. 1). Eyes were then placed on a shaker at room temperature, approximating the saccadic/translational motion to which living eyes would be subject, and ensuring that both aqueous and vitreal compartments were well-mixed, consistent with our mathematical modelling assumption (see the Model formulation section below). At times 20 min, 40 min, 1 hr and 3.5 hr, the aqueous and vitreous humour were extracted and processed to determine VEGF and ranibizumab concentrations (with *n* = 3 eyes per time point, per group).

**Figure 1:**
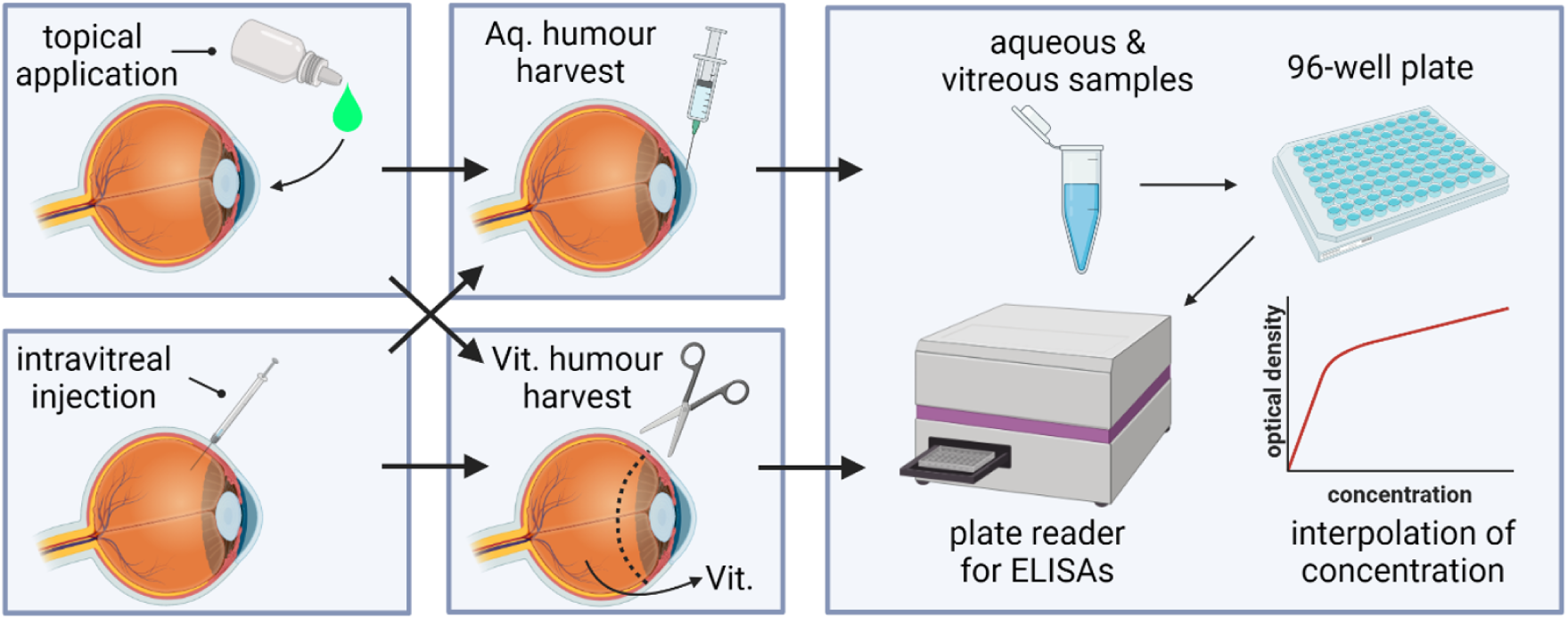
Diagram of the experimental procedure. Porcine eyes were treated either topically or via intravitreal injection. The aqueous humour was collected using a syringe, while the vitreous humour was harvested after removal of the anterior segment. Ranibizumab and VEGF levels were detected using ELISA, the optical density of the fluids measured using a plate reader and then converted into a concentration using the respective standard curves. (Figure created in https://BioRender.com.)

**Table 1:** Experimental treatment groups.

| Treatment | Delivery method | Num. eyes (*n*) |
| --- | --- | --- |
| Ranibizumab* and CPP<sup>†</sup> | Topical drop | 12 |
| Ranibizumab* and CPP<sup>†</sup> | Intravitreal injection | 12 |
| Ranibizumab* | Intravitreal injection | 12 |
| Sterile water | Intravitreal injection | 3 |

\*:Ranibizumab doses were all 1 mg ml<sup>-1</sup>.
<sup>†</sup>:CPP doses were all 100 mg ml<sup>-1</sup>.

#### Sample collection

Aqueous was extracted by making an incision in the cornea and using a Hamilton syringe to extract 200 µl of aqueous humour, which was stored in Eppendorfs on ice. To collect the vitreous, the cornea was surgically removed at the limbus and the vitreous dissected using tweezers, collected in a Falcon tube and homogenised. The homogenised vitreous was then centrifuged and the supernatant collected and stored at 4^◦^C (see Fig. 1).

### ELISA

The aqueous and vitreous samples were analysed by ELISA to detect levels of either VEGF-A (Invitrogen; Cat: #ES25RB) or ranibizumab (Abcam Cat: #ab282900). Both kits were performed according to the manufacturer’s instructions. Samples were diluted two-fold in Assay Buffer. Standards, controls, and samples (100 µl) were added to wells and incubated at room temperature for 30 min. After incubation, the plate was washed three times with wash buffer, then the (horseradish peroxidase) HRP-conjugate probe was added and incubated again. The incubation solution was discarded, and the wells were washed before adding (3,3’,5,5’-Tetramethylbenzidine) TMB Substrate, followed by a 10 min dark incubation. Stop solution was added, changing the colour from blue to yellow. A standard curve was prepared using provided standards, excluding standard zero. Optical density values were plotted against ranibizumab concentrations, and sample concentrations were obtained by multiplying by the dilution factor (see Fig. 1).

#### Statistical analysis

A two-sample Kolmogorov-Smirnov test was used for statistical analysis (to determine the likelihood that two sets of samples come from the same probability distribution), employing the Matlab (R2020a) routine kstest2 with default settings.

### Model formulation

We construct a mathematical model to predict the evolving concentrations of VEGF, ranibizumab and their compounds as they move and interact within the human or porcine eye. The model takes the form of a set of ordinary differential equations (ODEs), which describe the rate at which these concentrations change over time.

For simplicity, we do not model cell penetrating peptides (CPPs) explicitly. Whether they operate by binding to ranibizumab, or act directly on the cornea, the end result is to increase corneal permeability to ranibizumab, which is accounted for through an appropriate choice of the corneal permeability coefficient, β_Tear−Aq,r_ (cm hr^−1^; see Equations 3 and 5 below). In the case that CPPs do bind to ranibizumab, we assume that this has a negligible effect on its interaction with VEGF.

Biochemically, VEGF (V) can bind to either one or two molecules of ranibizumab (R), forming VEGF-ranibizumab (VR) and ranibizumab-VEGF-ranibizumab (RVR), respectively. ^26^ The reaction kinetics are as follows:

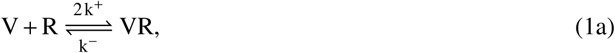

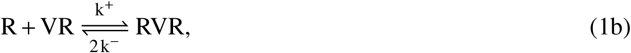

where, *k*^+^ (pmol^−1^ ml hr^−1^) is the binding rate of R to VR, and *k*^−^ (hr^−1^) is the unbinding rate of VR. We assume, with Hutton-Smith et al. (2016), ^17^ that the rate of binding(/unbinding) of R to(/from) each site on V is the same, regardless of whether the other site is filled. Since there are two empty binding sites available on V, the binding rate of R to V is 2*k*^+^, twice that of R to VR, where only one binding site is available. Similarly, the unbinding rate of RVR is 2*k*^−^, since either of the Rs can unbind to form R + VR.

The model is divided into 3 physical compartments: the tear film, the aqueous humour (filling the anterior segment) and the vitreous humour (filling the posterior segment). We describe the evolving concentrations of V, R, VR and RVR in the aqueous (Aq) and vitreal (Vit) compartments, and of R in the tear film (Tear) compartment, together with the variation in volume of the tear film with application and drainage of eye drops. See Table 2 for a full list of model variables, and Fig. 2 for a model diagram and schematic.

**Figure 2:**
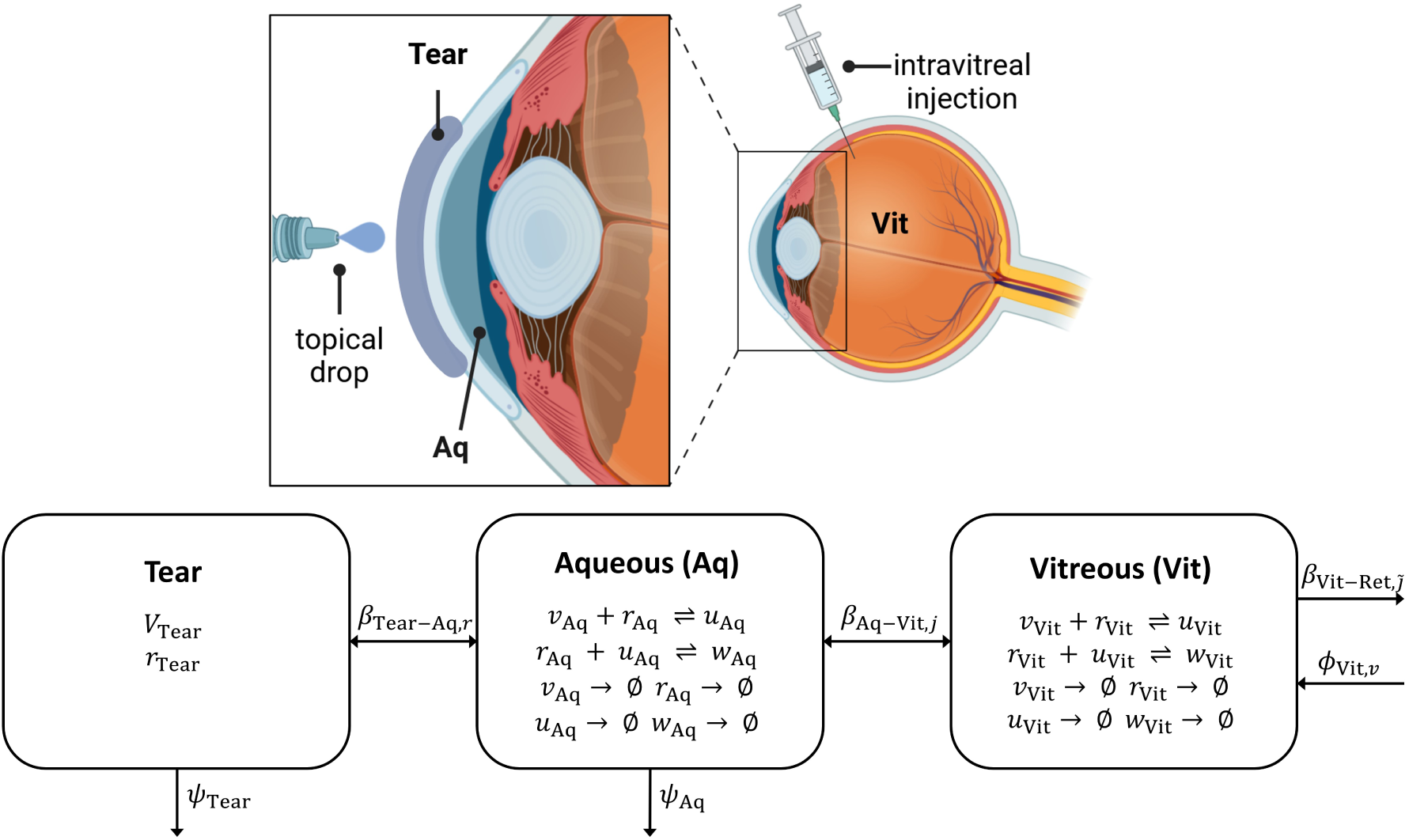
Anatomical and schematic diagrams for the mathematical model. **Top**: anatomical diagram, showing the locations of the tear film (Tear), aqueous humour (Aq) and vitreous humour (Vit), together with the application of topical and intravitreal treatments (the tear film and representations of treatment are not to scale). (Top panel created in https://BioRender.com.) **Bottom**: schematic diagram, summarising the movement of *v* (V), *r* (R), *u* (VR) and *w* (RVR) between compartments, their reactions (⇔) and degradation (→ ∅) within compartments, and the dynamic tear volume, *V*_Tear_ (see Table 2 for a full list of model variables). Parameters: β_Tear−Aq,r_ (cm hr^−1^), the permeability of the tear-aqueous interface to R; β_Aq−Vit,j_ (cm hr^−1^), for *j* ∈ {*v*, *r*, *u*, *w*}, the permeability of the aqueous-vitreous interface to V/R/VR/RVR; β_Vit−Ret_,^°^_j_ (cm hr^−1^), for ^°^*j r*, *u*, *w*, the permeability of the vitreo-retinal interface to R/VR/RVR; ψ_Tear_ (ml hr^−1^), the rate of fluid inflow/outflow to/from the tear film (causing dilution of R); ψ_Aq_ (ml hr^−1^), the rate of fluid inflow/outflow to/from the aqueous (causing dilution of V, R, VR and RVR); and φ_Vit,v_ (pmol hr^−1^), the rate at which the retina contributes V to the vitreous (see Table 3 for a full list of model parameters and their values).

**Table 2:** Independent and dependent variables used in the mathematical models (see Equations 2–13).

| Variable | Description (Units) |
| --- | --- |
| $t$ | Time (hr) |
| $V_{\text{Tear}}$ | Tear volume (ml) |
| $v_i$ | VEGF (V) concentration ($i \in \{\text{Aq, Vit}\}$) (pmol ml<sup>-1</sup>) |
| $r_i$ | Ranibizumab (R) concentration ($i \in \{\text{Tear, Aq, Vit}\}$) (pmol ml<sup>-1</sup>) |
| $u_i$ | VEGF-ranibizumab (VR) concentration ($i \in \{\text{Aq, Vit}\}$) (pmol ml<sup>-1</sup>) |
| $w_i$ | Ranibizumab-VEGF-ranibizumab (RVR) concentration ($i \in \{\text{Aq, Vit}\}$) (pmol ml<sup>-1</sup>) |

Time, $t$ , is the independent variable: a given for the models. The remaining variables are dependent variables: the quantities for which we are solving our models.

**Table 3:** Parameter values used in the mathematical models (see Equations 2–13), given to a maximum of three significant figures. Where a single value is given, this holds for both human and porcine eyes. Where two values are given, the first is for human eyes and the second is for porcine eyes. See Appendix A: Parameter justification for a description of how each parameter value was determined.

| Parameter | Description | Value | Source |
| --- | --- | --- | --- |
| $\tau_{\text{loss}}$ | Time for tear volume to return to normal (human) / zero (porcine) | $5.00 \times 10^{-2}$ hr<br>$5.00 \times 10^{-2}$ hr | 37–41<br>this study and 37–41 |
| $V_{\text{Drop}}$ | Volume of eye drop | $4.50 \times 10^{-2}$ ml | this study |
| $V_{\text{TearNorm}}$ | Volume of normal tear film | $6.35 \times 10^{-3}$ ml | 42, 43 |
|  |  | N/A | — |
| $V_{\text{TearRes}}$ | Volume of tear reservoir | $3.00 \times 10^{-2}$ ml | 38, 40, 44, 45 |
|  |  | N/A | — |
| $V_{\text{Aq}}$ | Volume of aqueous | 0.160 ml | 46 |
|  |  | 0.310 ml | 47, 48 |
| $V_{\text{Vit}}$ | Volume of vitreous | 4.50 ml | 17, 47, 49, 50 |
|  |  | 3.10 ml | 47, 48, 51, 52 |
| $A_{\text{Tear-Aq}}$ | Area of the tear-aqueous interface | 1.30 cm<sup>2</sup> | calc. from data in 53, 54 |
|  |  | 1.51 cm<sup>2</sup> | calc. from data in this study and 55 |
| $A_{\text{Aq-Vit}}$ | Area of the aqueous-vitreous interface | 0.349 cm<sup>2</sup> | calc. from data in 56 |
|  |  | 0.301 cm<sup>2</sup> | calc. from data in 53, 55, 57–62 |
| $A_{\text{Vit-Ret}}$ | Area of the vitreo-retinal interface | 10.9 cm<sup>2</sup> | calc. from data in 63 |
|  |  | N/A | — |
| $k^+$ | Binding rate of R to VR (to form RVR) | $0.576 \text{ pmol}^{-1} \text{ ml hr}^{-1}$ | 64 |
| $k^-$ | Unbinding rate of VR (to form V + R) | $2.63 \times 10^{-2} \text{ hr}^{-1}$ | 64 |
| $\delta_{i,j}$ | Rate of molecular degradation ($i \in \{\text{Aq, Vit}\}$ and $j \in \{v, r, u, w\}$) | $0 \text{ hr}^{-1}$ | simplifying assumption |
| $\beta_{\text{Tear-Aq},r}$ | Permeability of the tear-aqueous interface to R | $1.10 \times 10^{-6} \text{ cm hr}^{-1}$ | inferred from porcine value using data in 53, 57, 62, 65 |
| | | $5.93 \times 10^{-7} \text{ cm hr}^{-1}$ | model fit |
| $\beta_{\text{Aq-Vit},v}$ | Permeability of the aqueous-vitreous interface to V | $9.85 \times 10^{-1} \text{ cm hr}^{-1}$ | inferred from $\beta_{\text{Aq-Vit},r}$ using data in 17, 66, 67 |
| $\beta_{\text{Aq-Vit},r}$ | Permeability of the aqueous-vitreous interface to R | $9.29 \times 10^{-1} \text{ cm hr}^{-1}$ | model fit |
| $\beta_{\text{Aq-Vit},u}$ | Permeability of the aqueous-vitreous interface to VR | $7.55 \times 10^{-1} \text{ cm hr}^{-1}$ | inferred from $\beta_{\text{Aq-Vit},r}$ using data in 17, 66, 67 |
| $\beta_{\text{Aq-Vit},w}$ | Permeability of the aqueous-vitreous interface to RVR | $6.53 \times 10^{-1} \text{ cm hr}^{-1}$ | inferred from $\beta_{\text{Aq-Vit},r}$ using data in 17, 66, 67 |
| $\beta_{\text{Vit-Ret},r}$ | Permeability of the vitreo-retinal interface to R | $6.80 \times 10^{-4} \text{ cm hr}^{-1}$ | 68 |
| | | $0 \text{ cm hr}^{-1}$ | — |
| $\beta_{\text{Vit-Ret},u}$ | Permeability of the vitreo-retinal interface to VR | $6.44 \times 10^{-4} \text{ cm hr}^{-1}$ | 68 |
| | | $0 \text{ cm hr}^{-1}$ | — |
| $\beta_{\text{Vit-Ret},w}$ | Permeability of the vitreo-retinal interface to RVR | $6.23 \times 10^{-4} \text{ cm hr}^{-1}$ | 68 |
| | | $0 \text{ cm hr}^{-1}$ | — |
| $\phi_{\text{Vit},v}$ | Rate at which retina contributes V to vitreous | $2.34 \times 10^{-4} \text{ pmol hr}^{-1}$ | mean value from 17 |
| | | $0 \text{ pmol hr}^{-1}$ | — |
| $\psi_{\text{Tear}}$ | Rate of inflow/outflow to/from tear film | $7.20 \times 10^{-2} \text{ ml hr}^{-1}$ | 42 |
| | | $0 \text{ ml hr}^{-1}$ | — |
| $\psi_{\text{Aq}}$ | Rate of inflow/outflow to/from aqueous | $0.150 \text{ ml hr}^{-1}$ | 17, 65, 69, 70 |
| | | $0 \text{ ml hr}^{-1}$ | — |
| $v_{\text{Aq},\text{init}}$ | Initial V concentration in aqueous | $1.56 \times 10^{-3} \text{ pmol ml}^{-1}$ | model steady-state without R |
| | | $0 \text{ pmol ml}^{-1}$ | simplifying assumption / this study |
| $v_{\text{Vit},\text{init}}$ | Initial V concentration in vitreous | $2.24 \times 10^{-3} \text{ pmol ml}^{-1}$ | model steady-state without R |
| | | $0 \text{ pmol ml}^{-1}$ | simplifying assumption / this study |
| $r_{\text{Dose}}$ | R dose concentration | $2.07 \times 10^4 \text{ pmol ml}^{-1}$ | this study and 17, 66 |
| $r_{\text{Tear},\text{init}}$ | Initial R concentration in tear | $1.81 \times 10^4 \text{ pmol ml}^{-1}$ | calculated |
| | | $2.07 \times 10^4 \text{ pmol ml}^{-1}$ | calculated |
| $r_{\text{Vit},\text{init}}$ | Initial R concentration in vitreous | $2.07 \times 10^2 \text{ pmol ml}^{-1}$ | calculated |
| | | $4.29 \text{ pmol ml}^{-1}$ | this study |

Parameters are fixed quantities which do not change during a simulation.

In choosing an ODE model, we are making the assumption that the chemical species modelled in each physical compartment are well-mixed. This is a justified simplification given the low volume of the tear film and aqueous, and the mixing effect of the fluid flow within them, and given the mixing effect within the vitreous of saccadic eye rotations, especially in the partially liquefied aged vitreous of AMD patients. ^27–31^

In what follows, we define the model in its most general form, which applies to an *in vivo* human eye with topical application of ranibizumab. We then describe how the equations are simplified for intravitreal injections, for the untreated case, and/or for an *ex vivo* porcine eye. Two subcases of the topical treatment case are considered: *drop-based therapy* (as in our experiments; see Experimental methods section) and *drug-eluting contact lenses*. ^10^ For topical drops, tear volume and ranibizumab concentration deplete following drop application, while for drug-eluting contact lenses, tear volume and ranibizumab concentration are assumed to remain constant while a lens is worn. (When fitting to *ex vivo* porcine experimental data, we also consider the scenario where the tear ranibizumab concentration depletes, but the tear/applied drop volume is held constant.)

The Matlab (R2020a) routine ode45, which employs an explicit Runge-Kutta method, was used for solving all versions of our ODE model.

#### Tear film (Tear) compartment

This compartment is used only when considering topical treatments. When using a drug-eluting contact lens, the tear film volume, *V*_Tear_(*t*) (ml), remains fixed at its normal volume, *V*_TearNorm_ = 6.35 × 10^−3^ ml; whereas, when treating with eye drops, *V*_Tear_ is a function of time, *t* (hr), since the last drop was applied. The eye drops in our study have a volume of *V*_Drop_ = 4.5 × 10^−2^ ml, while the maximum volume of fluid that can be held in the tear film, known as the reservoir volume, is *V*_TearRes_ = 3 × 10^−2^ ml. Therefore, following the application of an eye drop, the tear volume increases to *V*_TearRes_, while the remaining liquid (with a volume equal to *V*_TearNorm_ + *V*_Drop_ − *V*_TearRes_ = 2.14 × 10^−2^ ml) flows immediately off of the eye. While *V*_Tear_(*t*) > *V*_TearNorm_, the rate of tear drainage exceeds that of the inflow, such that *V*_Tear_(*t*) returns to *V*_TearNorm_ in time τ_loss_ (hr), where we assume that volume reduction proceeds linearly. Thus, we have the following algebraic equations for tear film volume:

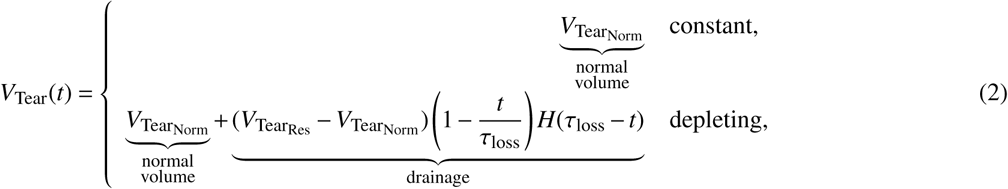

where, *H*(·) is the Heaviside step function, defined as

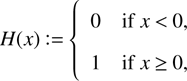

which is required to prevent *V*_Tear_(*t*) going below *V*_TearNorm_ when *t* > τ_loss_.

The tear film ranibizumab concentration, *r*_Tear_(*t*) (pmol ml^−1^), is assumed to be fixed in the drug-eluting contact lens case, such that the rate of change of *r*_Tear_ over time, 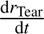, is zero. While, in the drop-based therapy case, *r*_Tear_ decreases over time due to its diffusion across the cornea into the aqueous, and due to dilution of the tear film as new fluid is added and old fluid is drained away (ranibizumab is assumed to be too large and hydrophilic to pass across the conjunctiva). VEGF concentrations in the human tear film have been measured to lie in the range 7.48 × 10^−4^–3.08 × 10^−1^ pmol ml^−1^ across healthy and AMD patients, ^32–34^ around five to seven orders of magnitude smaller than the tear ranibizumab concentration following topical application, *r*_Tearinit_ = 1.81 × 10^4^ pmol ml^−1^ (see Appendix A: Parameter justification). Therefore, we neglect VEGF and its compounds (V, VR and RVR) in the tear film. We assume that CPPs aid only the passage of ranibizumab, and only then in passing from the tear film to the aqueous. Therefore, we assume that V, VR and RVR cannot pass across the cornea, and that the flux of ranibizumab across the cornea depends only upon its concentration in the tear film and not upon its concentration in the aqueous. Given that the volume of the tear film (*V*_TearNorm_ = 6.35 × 10^−3^ ml) is over an order of magnitude smaller than that of the aqueous (*V*_Aq_ = 0.16 ml), we anticipate that the effect of the inclusion of these additional fluxes upon aqueous chemical concentrations would be negligible in any case. Thus, we have the following equation:

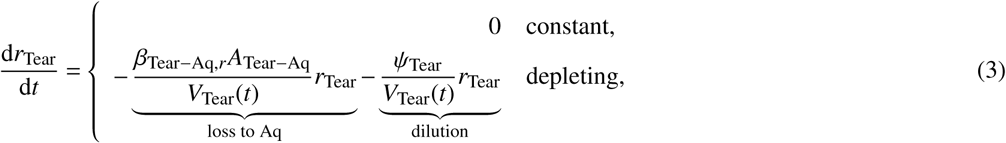

where β_Tear−Aq,_*_r_* (cm hr^−1^) is the permeability of the cornea to ranibizumab (in the presence if CPPs), ψ_Tear_ (ml hr^−1^) is both the rate of tear film production and the rate of tear film drainage (when *V*_Tear_(*t*) = *V*_TearNorm_), and *A*_Tear−Aq_ (cm^2^) is the corneal area (that is, the area of the Tear-Aq interface).

#### Aqueous (Aq) compartment

We describe the evolving concentrations of all 4 biomolecules in the aqueous: V, *v*_Aq_(*t*) (pmol ml^−1^), R, *r*_Aq_(*t*) (pmol ml^−1^), VR, *u*_Aq_(*t*) (pmol ml^−1^), and RVR, *w*_Aq_(*t*) (pmol ml^−1^). The dynamics of these biomolecules are governed by their reaction kinetics (Equations 1a–1b) and the Law of Mass Action (which states that the rate of a reaction is directly proportional to the product of the concentrations of the reactants), ^35^ their decay(/biological degradation), influx from the tear film (through the cornea; R only), exchange with the vitreous (through the suspensory ligaments / zonule, via diffusive flux), and dilution (through production and drainage of the aqueous humour). (Current evidence suggests that aqueous fluid may flow posteriorly, into the vitreous. ^36^ Given that the rate of flow has yet to be determined, ^36^ we neglect it here, noting that its inclusion would both increase delivery of topically applied ranibizumab to the posterior segment and reduce ranibizumab loss from the vitreous following intravitreal injection. Ranibizumab is assumed to be too large and hydrophilic to pass across the blood vessel endothelia in the iris and ciliary body.) This gives rise to the following equations:

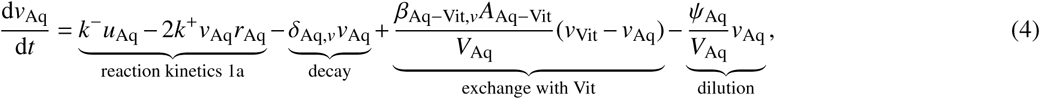

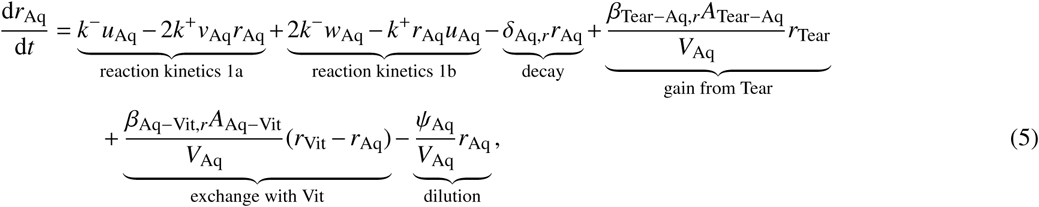

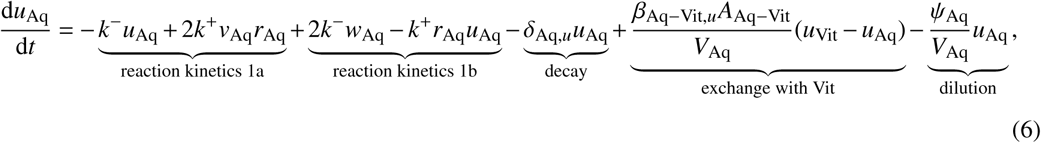

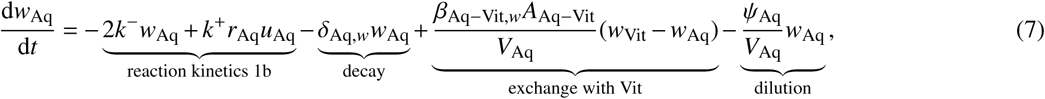

where δ_Aq,_ *_j_* (hr^−1^) is the rate of molecular degradation of *j* ∈ {*v*, *r*, *u*, *w*} in the aqueous, β_Aq−Vit,j_ (cm hr^−1^) is the permeability of the aqueous-vitreous interface (suspensory ligaments) to *j* ∈ {*v*, *r*, *u*, *w*}, ψ_Aq_ (ml hr^−1^) is the rate of inflow(/outflow) of aqueous humour to(/from) the anterior segment, *V*_Aq_ (ml) is the aqueous volume, and *A*_Aq−Vit_ (cm^2^) is the area of the aqueous-vitreous interface.

#### Vitreal (Vit) compartment

We describe the evolving concentrations of all 4 biomolecules in the vitreous: V, *v*_Vit_(*t*) (pmol ml^−1^), R, *r*_Vit_(*t*) (pmol ml^−1^), VR, *u*_Vit_(*t*) (pmol ml^−1^), and RVR, *w*_Vit_(*t*) (pmol ml^−1^). As in the aqueous, the dynamics of these biomolecules are governed by their reaction kinetics, their decay(/biological degradation), and their exchange between the aqueous and vitreous. In addition, R, VR and RVR are lost to the retinal/choroidal tissue via diffusive flux, and V is gained from the retina, where it is produced by the retinal pigment epithelium (RPE). Consequently, we have the following equations:

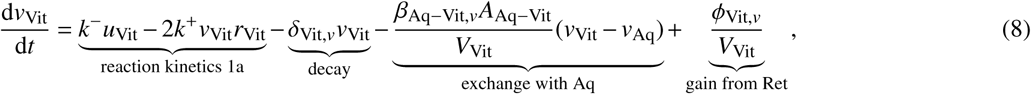

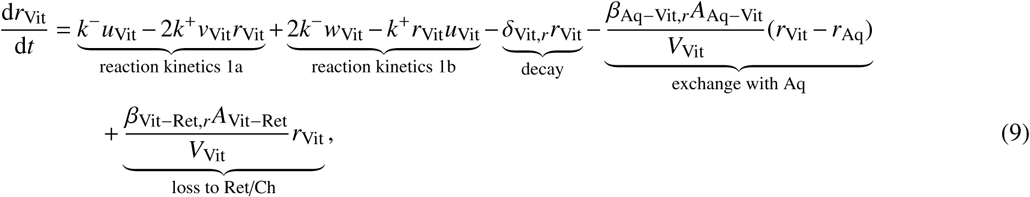

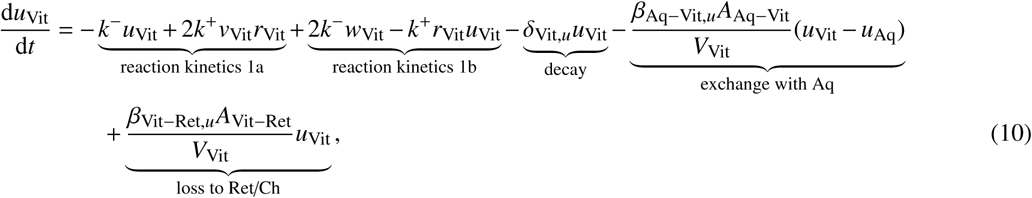

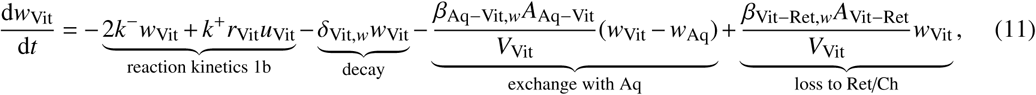

where δ_Vit,_ *_j_* (hr^−1^) is the rate of molecular degradation of *j* ∈ {*v*, *r*, *u*, *w*} in the vitreous, β_Vit−Ret,j_ (cm hr^−1^) is the permeability of the vitreo-retinal interface (inner limiting membrane, ILM) to *j* ∈ {*r*, *u*, *w*}, φ_Vit,v_ (pmol hr^−1^) is the rate at which the retina contributes VEGF (produced by the RPE) to the vitreous, *V*_Vit_ (ml) is the vitreal volume, and *A*_Vit−Ret_ (cm^2^) is the area of the vitreo-retinal interface (ILM). The loss to the retina/choroid terms are taken to depend only upon the R/VR/RVR concentration in the vitreous since the choroidal blood flow quickly removes these species.

#### Initial conditions

In addition to the governing equations given above (Equations 2–11), we impose the following initial conditions to fully define our model:

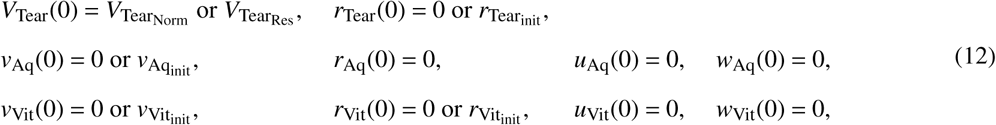

where *r*_Tearinit_ (pmol ml^−1^), *v*_Aqinit_ (pmol ml^−1^), *v*_Vitinit_ (pmol ml^−1^) and *r*_Vitinit_ (pmol ml^−1^) are positive constants. In the case where an eye drop is applied at *t* = 0 hr, the initial tear volume is *V*_TearRes_, otherwise *V*_Tear_(0) = *V*_TearNorm_ (technically, the tear volume does not require an initial condition, since it defined by an algebraic equation; however, we include it here for completeness and clarity). For an initial topical treatment (drops or contact lenses), the initial tear ranibizumab concentration *r*_Tear_(0) = *r*_Tearinit_, while without topical treatment *r*_Tear_(0) = 0. Similarly, for an initial intravitreal treatment, the initial vitreal ranibizumab concentration is *r*_Vitinit_, while without intravitreal treatment, *r*_Vit_(0) = 0. Treatment is never applied directly to the aqueous, hence *r*_Aq_(0) = 0. The initial aqueous and vitreal VEGF concentrations, *v*_Aqinit_ and *v*_Vitinit_, are chosen to be the steady-state VEGF concentrations in the absence of ranibizumab treatment. Finally, the initial concentrations of VR and RVR are chosen to be zero, since ranibizumab has not yet had an opportunity to interact with VEGF. See Table 3 for a full list of model parameters (fixed quantities) and Appendix A: Parameter justification for a description of how each parameter value was determined.

#### Cases and submodels

In what follows, we explain how our mathematical model differs between the *in vivo* human eye and the *ex vivo* porcine eye, and under each treatment condition, for scenarios in which at most a single dose is applied (at *t* = 0 hr; see Repeated dosing section below for repeated dosing scenarios).

• *In Vivo* Human Eye

**–** Topical Treatment

∗ Drug-eluting contact lens: model is as stated in Equations 2–12, taking the first option in Equations 2 and 3, such that *V*_Tear_ = *V*_TearNorm_ and *r*_Tear_ = *r*_Tearinit_ are constants, and choosing *v*_Aq_(0) = *v*_Aqinit_, *v*_Vit_(0) = *v*_Vitinit_ and *r*_Vit_(0) = 0.

∗ Drop-based therapy: model is as stated in Equations 2–12, taking the second option in Equations 2 and 3, and choosing *V*_Tear_(0) = *V*_TearRes_, *r*_Tear_(0) = *r*_Tearinit_, *v*_Aq_(0) = *v*_Aqinit_, *v*_Vit_(0) = *v*_Vitinit_ and *r*_Vit_(0) = 0.

**–** Intravitreal injection: equations concerned with the tear film are removed (Equations 2 and 3), together with their associated dependent variables (*V*_Tear_ and *r*_Tear_) and initial conditions (Equation 12, top line), as is the gain from tear film term from Equation 5, while *v*_Aq_(0) = *v*_Aqinit_, *v*_Vit_(0) = *v*_Vitinit_ and *r*_Vit_(0) = *r*_Vitinit_.

**–** No treatment: this case is considered only for determining the untreated steady-state values of *v*_Aq_ and *v*_Vit_. These values are used as the initial VEGF values (*v*_Aqinit_ and *v*_Vitinit_) in all other *in vivo* human eye simulations. Only the VEGF equations (Equations 4 and 8) and their associated dependent variables (*v*_Aq_ and *v*_Vit_) are used, together with the initial conditions *v*_Aq_(0) = 0 and *v*_Vit_(0) = 0. All terms in Equations 4 and 8 involving ranibizumab (R) or its compounds with VEGF (VR and RVR) are removed.

• *Ex Vivo* Porcine Eye

**–** Topical Treatment

∗ Drug-eluting contact lens: we do not consider this scenario for the *ex vivo* porcine eye since we do not have experimental data for this treatment type.

∗ Drop-based therapy: the model is similar to that for the *in vivo* human eye (above), but with the following modifications — we replace Equation 2 with Equation 13 (below), and *V*_Tear_(0) = *V*_Drop_. We also modify the parameters (see Table 3); in particular, ψ_Tear_ = 0, ψ_Aq_ = 0, φ_Vit,v_ = 0, and β_Vit−Ret,j_ = 0 for *j* ∈ {*r*, *u*, *w*}, since there is no tear or aqueous production/drainage, and no VEGF production or choroidal blood flow in an *ex vivo* eye.

**–** Intravitreal injection: the model is the same as for the *in vivo* human eye (above), but with modified parameters (see Table 3). In particular, ψ_Aq_ = 0, φ_Vit,v_ = 0, and β_Vit−Ret,j_ = 0 for *j* ∈ {*r*, *u*, *w*}, since there is no aqueous production/drainage, VEGF production or choroidal blood flow in an *ex vivo* eye.

**–** No treatment: this scenario is not considered for the *ex vivo* porcine eye, since we use values for *v*_Aqinit_ and *v*_Vitinit_ based on our experimental measurements in this case, rather than the steady-state solution to the untreated problem, as we do for the *in vivo* human eye (see above).

We consider two forms for the tear volume equation in the *ex vivo* porcine eye: a *depleting volume* case in which the drop flows off of the cornea in time τ_loss_ (linearly, as with the *in vivo* human eye), and the limiting, *constant volume* case, where the full drop remains on the cornea throughout the experiment. This gives rise to the following equations:

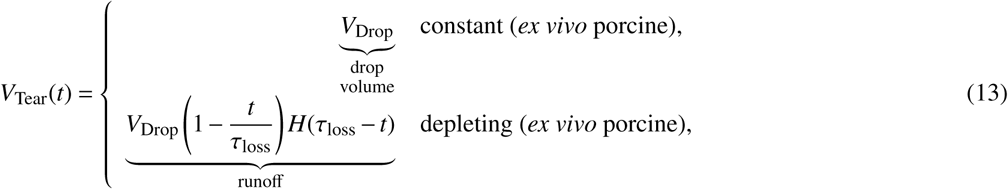

where the full drop volume is lost in the volume loss case since a tear film cannot be maintained in the absence of the surrounding tissues. We consider both constant volume and volume loss cases since it is impractical to precisely characterise the time for a drop to flow off of the cornea (visual observation suggests that the majority of the drop volume remains on the cornea during the experiment), and to allow us to assess the influence of the rate of drop volume loss when fitting other parameters to the experimental data (see Model fitting below).

#### Repeated dosing

Alongside the single dose scenario, we also consider repeated dosing for drop, contact lens and intravitreal treatments in the *in vivo* human eye. In this case, the initial conditions given in Equation 12 hold only for the initial time interval (that is, the interval before the second dose is applied if the first is applied at *t* = 0 hr, or the interval before the first dose is applied otherwise). For each dose applied after *t* = 0 hr, we solve the governing equations (2–11) up to the time at which the new dose is applied, *t* = *t*_dose_*_n_* hr say, at which point we modify the relevant dependent variables to account for the new treatment, which provides the initial conditions for the next time interval, *t* ∈ (*t*_dose_*_n_*, *t*_dose_*_n_*_+1_].

Upon the application of a drop, *V*_Tear_(*t*) jumps from *V*_TearNorm_ to *V*_TearRes_, with the excess volume (*V*_TearNorm_ + *V*_Drop_ − *V*_TearRes_) flowing immediately off of the eye. With the exception of *r*_Tear_ and/or *r*_Vit_, the initial conditions for each biochemical dependent variable are equal to their values at the final time point in the previous treatment interval. Upon application of a drop/contact lens and/or intravitreal treatment, the initial conditions for *r*_Tear_ and/or *r*_Vit_ respectively will be discontinuous with their values at the final time point in the previous treatment interval, with new values calculated as follows.

When a new eye drop is applied, the fluid in the drop immediately merges and mixes with the fluid in the tear film, such that 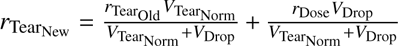, where *r*_Tear_ (pmol ml^−1^) is the tear film ranibizumab concentration immediately after the application of a drop, *r*_TearOld_ (pmol ml^−1^) is the tear film ranibizumab concentration immediately before the application of a drop, and *r*_Dose_ (pmol ml^−1^) is the ranibizumab concentration within the drop, prior to application. Thus, the mixing is assumed to occur sufficient rapidly that the ranibizumab concentration equilibrates throughout the tear film/drop, prior to the excess volume (*V*_TearNorm_ + *V*_Drop_ − *V*_TearRes_) leaving the tear film. If the previous treatment at *t*_dose_*_n_*_−1_ was also in the form of an eye drop, then we choose *t*_dose_*_n_* such that *t*_dose_*_n_* − *t*_dose_*_n_*_−1_ ≥ τ_loss_, ensuring that *V*_Tear_(*t*) has returned to *V*_TearNorm_ when the new drop is applied.

When a new drug-eluting contact lens is inserted, it is assumed that the tear ranibizumab concentration immediately takes a value of *r*_Tearinit_, and that this is maintained throughout the period for which it is worn. Upon removal of the contact lens, the tear ranibizumab concentration decays as described by the depleting form of Equation 3.

When a new intravitreal injection is administered, it is assumed that the ranibizumab concentration quickly equilibrates throughout the vitreous, such that 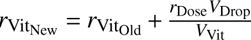, where *r*_VitNew_ (pmol ml) is the vitreal ranibizumab concentration immediately after the application of an injection, and *r*_VitOld_ (pmol ml^−1^) is the vitreal ranibizumab concentration immediately before the application of an injection (the injected volume is the same as the drop volume, hence we use the same parameter, *V*_Drop_, for both). It is assumed that the injected volume does not affect the vitreal volume, the additional volume being negligible (given that *V*_Drop_ « *V*_Vit_) and quickly drained.

We also consider treatment regimens which combine topical (drop and contact lens) and intravitreal doses. In this case, each treatment type follows the rules described above.

In the case where a single dose is applied at *t* = 0 hr, time, *t*, can simply be measured from *t* = 0 hr. The situation is a little more complicated in the case of repeated dosing, since solution of the governing equations (2–11) must be halted and re-initialised for every new dose. As such, it is helpful to distinguish between what we shall term ‘local’ and ‘global’ time. Global time spans the full duration of a simulation, and is the time plotted in figures, while local time is the time ‘seen’ by the governing equations, and is reset to zero upon the application of each treatment. For notational simplicity, we use *t* to denote both local and global time, with the local/global distinction being understood from the context in which it is used.

## Results

### Porcine eyes

#### Experimental results

Control ranibizumab measurements in the aqueous were 0 pmol ml^−1^, while those in the vitreous were 0.348±0.459 pmol ml^−1^, despite ranibizumab being absent. This is likely due to the effects of vitreal proteins being mis-detected as ranibizumab. Control VEGF measurements in the aqueous were 9.87×10^−3^±1.24×10^−3^ pmol ml^−1^, while those in the vitreous were 0 pmol ml^−1^. It is unclear why VEGF was not detected in the vitreal controls. Given that VEGF levels in the vitreous for topical therapy (for which vitreal ranibizumab levels are within the range of controls) are at similar levels to the aqueous control values, we assume that vitreal control values should be the roughly the same as aqueous control values.

Three treatments were tested: 1. topical treatment with ranibizumab and CPP, 2. intravitreal treatment with ranibizumab and CPP, and 3. intravitreal treatment with ranibizumab alone (see Table 4 for a summary of the experimental results).

**Table 4:**
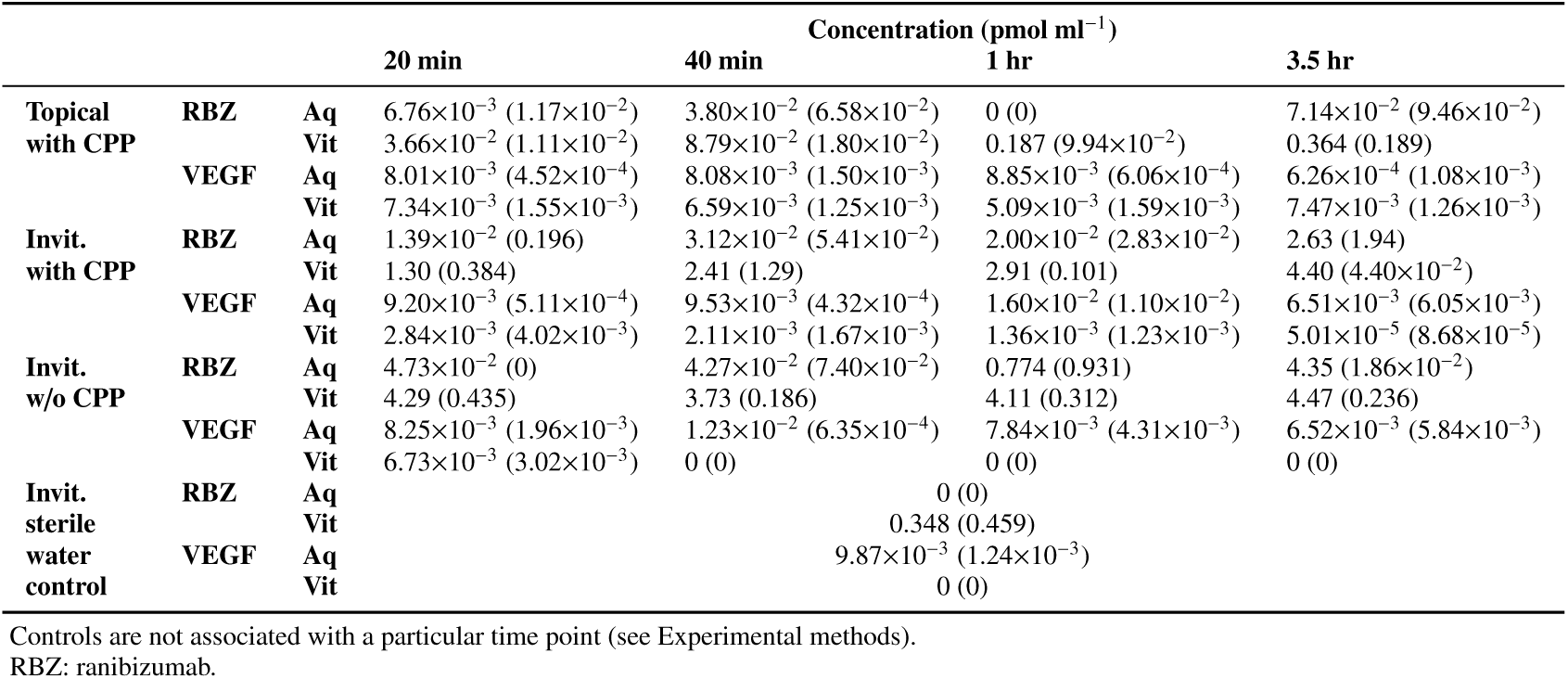
Experimental results for treatment of *ex vivo* porcine eyes. Stated values are means, with standard deviations in parentheses (*n* = 3 for each data point). All values are given to three significant figures. These data are plotted in Fig. 3.

First, for topical treatment with ranibizumab and CPP (Fig. 3 (top row)), the aqueous ranibizumab levels increased significantly, vs. controls, by 3.5 hr (7.14×10^−2^±9.46×10^−2^ pmol ml^−1^; *p* = 0.03), coincident with a significant (*p* = 0.03) drop in aqueous VEGF between 1 hr (8.85×10^−3^±6.06×10^−4^ pmol ml^−1^) and 3.5 hr (6.26×10^−4^±1.08×10^−3^ pmol ml^−1^). Vitreal ranibizumab increased gradually, with significant (*p* = 0.03) increase between 40 min (8.79×10^−2^±1.80×10^−2^ pmol ml^−1^) and 3.5 hr (0.364±0.189 pmol ml^−1^); though remaining within the range of the controls (0.348±0.459 pmol ml^−1^), while vitreal VEFG remained constant. The aqueous ranibizumab measurements at 1 hr were all 0 pmol ml^−1^, this was somewhat anomalous given that ranibizumab was detected at all other time points, thus we neglect this point in the Model fitting section below.

**Figure 3:**
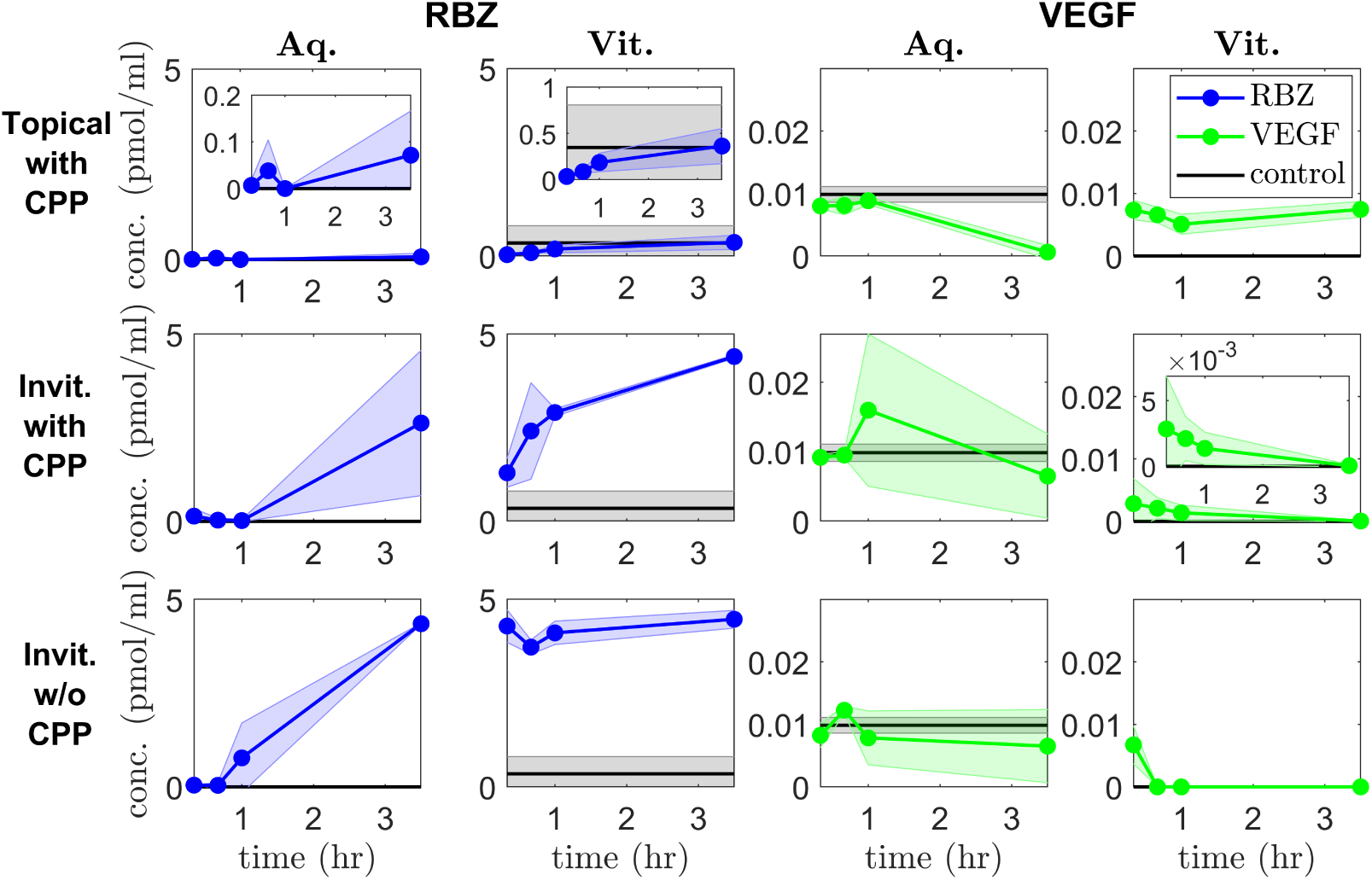
Experimental results for treatment of *ex vivo* porcine eyes. Panels show ranibizumab (RBZ, **first two columns**) and VEGF (**last two columns**) concentrations in the aqueous (Aq, **columns 1 and 3**) and vitreous (Vit, **columns 2 and 4**), at *t* = 20 min, 40 min, 1 hr and 3.5 hr. **Top row**: topical treatment (45 µl) with ranibizumab (1 mg ml^−1^) and CPP (100 mg ml^−1^). Ranibizumab is detected in significant quantities in the aqueous and appears to enter the vitreous, though it only has a significant effect in reducing aqueous VEGF levels, leaving vitreal VEGF levels unaffected. **Middle row**: intravitreal (invit.) treatment (45 µl) with ranibizumab (1 mg ml^−1^) and CPP (100 mg ml^−1^). As expected, vitreal ranibizumab levels are high, resulting in a significant reduction in vitreal VEGF levels. **Bottom row**: intravitreal treatment (45 µl) with ranibizumab only (1 mg ml^−1^). Interestingly, intravitreal treatment is more effective in reducing vitreal VEGF in the absence of CPPs. See Table 4 for numerical values of data points.

Second, for intravitreal treatment with ranibizumab and CPP (Fig. 3 (middle row)), aqueous ranibizumab was first detected in significant quantities, compared to controls, at 3.5 hr (2.63±1.94 pmol ml^−1^; *p* = 0.03), while aqueous VEFG remained constant. Vitreal ranibizumab increased steadily and significantly (*p* = 0.03) from 20 min (1.30±0.384 pmol ml^−1^) to 3.5 hr (4.40±4.40×10^−2^ pmol ml^−1^), coincident with a significant (*p* = 0.03) drop in vitreal VEGF from 40 min (2.11×10^−3^±1.67×10^−3^ pmol ml^−1^) to 3.5 hr (5.01×10^−5^±8.68×10^−5^ pmol ml^−1^).

Third, for intravitreal treatment with ranibizumab alone (Fig. 3 (bottom row)), aqueous ranibizumab was present in significant quantities, vs. controls, by 1 hr (0.774±0.931 pmol ml^−1^; *p* = 0.03), while aqueous VEFG remained constant. Vitreal ranibizumab maintained constant values (∼4 pmol ml^−1^) for all time points, while vitreal VEGF was undetectable from 40 min onwards.

Detected vitreal ranibizumab values following intravitreal ranibizumab injections (with and without CPP) are much lower than would be anticipated based on the injected dose concentration (and volume). Given the injected dose, an initial intravitreal ranibizumab concentration of 3.00 × 10^2^ pmol ml^−1^ would be expected (see Appendix A: Parameter justification); however, measured vitreal values do not exceed 5 pmol ml^−1^. It may be that, despite being placed on a shaker, vitreal ranibizumab does not spread out uniformly from the injection site over the time-frame of our experiments, and that our samples were taken from a portion of the vitreous away from the injection site.

### Model fitting

All *ex vivo* porcine parameter values were taken or calculated from the literature, or taken directly from our experimental data, except for the permeabilities of the Tear-Aq and Aq-Vit interfaces to ranibizumab, β_Tear−Aq,_*_r_* and β_Aq−Vit,_*_r_* respectively, which were determined by fitting our mathematical model to the experimental data.

Data from topical drop administration with CPP experiments were used for fitting, for which a single drop is applied at *t* = 0 hr. Equations 3–13 were solved with depleting tear ranibizumab concentration, constant tear volume and in the absence of VEGF. Simulations were initialised at *t* = 0 hr with *r*_Tear_(0) = *r*_Dose_ = 2.07 × 10^4^ pmol ml^−1^, *r*_Aq_(0) = 0 pmol ml^−1^ and *r*_Vit_(0) = 0 pmol ml^−1^, and the Matlab (R2020a) routine fminsearch (which uses a Nelder-Mead simplex method) applied (with default settings) to minimise the mean squared error between the data and the model predictions. Fitting was performed for *r*_Aq_ at *t* = 20 min, 40 min and 3.5 hr to the mean data points, which were deemed to be more accurate than the vitreal ranibizumab measurements, and neglecting the aqueous data point at *t* = 1 hr, which appears anomalous (see Experimental results).

A good fit was achieved in the aqueous (*model fit 1*: β_Tear−Aq,*r*_ = 5.93 × 10^−7^ cm hr^−1^ and β_Aq−Vit,*r*_ = 0.929 cm hr^−1^); however, it is not possible to find a good fit in both aqueous and vitreal compartments simultaneously, since measured vitreal ranibizumab concentrations are higher than those in the aqueous, and ranibizumab must move down a concentration gradient from the aqueous to the vitreous for topical treatment (see the top row of Fig. 4).

**Figure 4:**
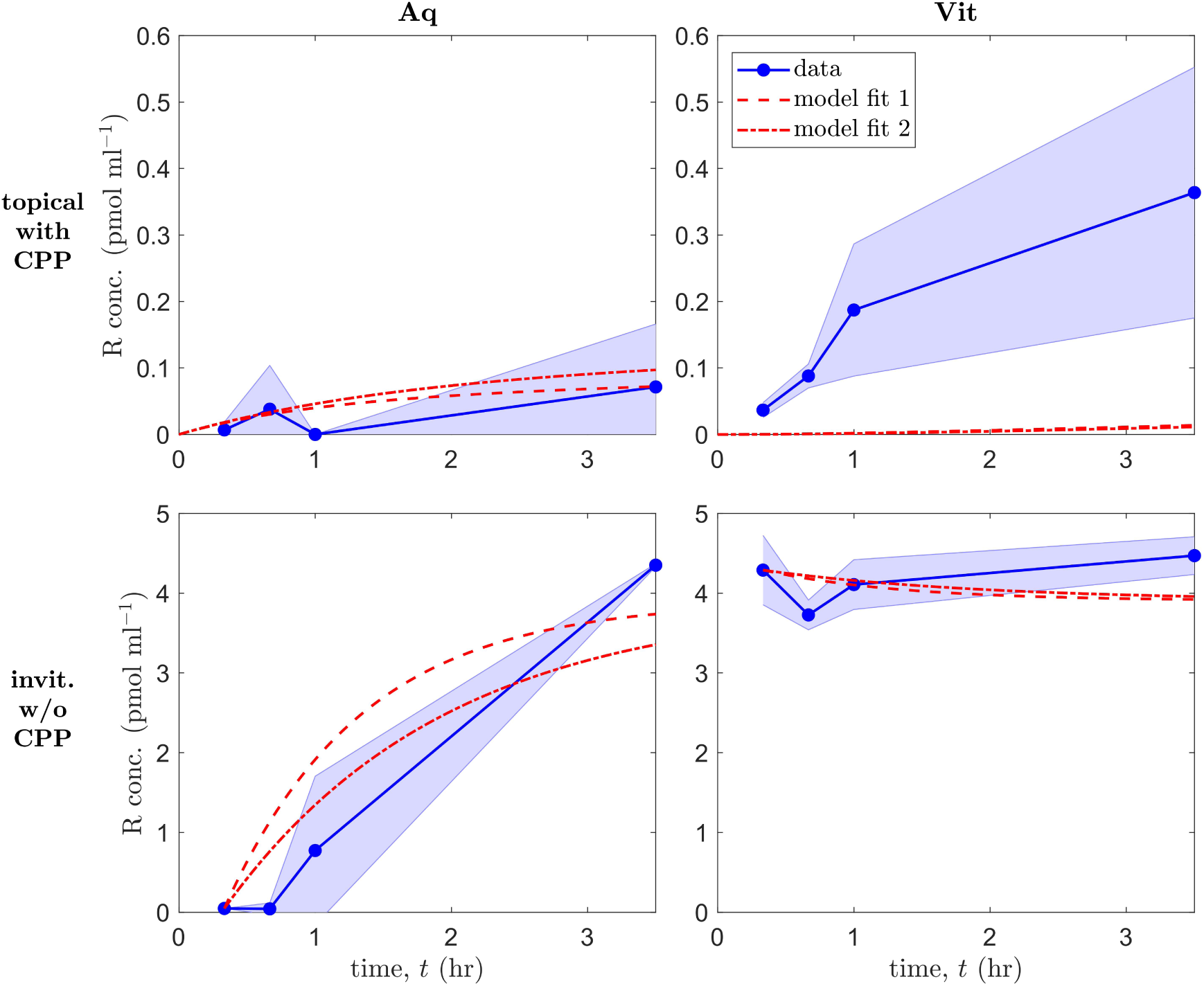
Mathematical model fits to *ex vivo* porcine data. Panels show ranibizumab (R) concentrations in the aqueous (Aq, **left column**) and vitreous (Vit, **right column**), for topical drop administration with CPPs (**top row**) and for intravitreal injection without CPPs (**bottom row**). **Top row**: a single drop is applied at *t* = 0 hr; simulations start at *t* = 0 hr with *r*_Tear_(0) = *r*_Dose_ = 2.07 10^4^ pmol ml^−1^, *r*_Aq_(0) = 0 pmol ml^−1^ and *r*_Vit_(0) = 0 pmol ml^−1^; fitting was performed for *r*_Aq_ at *t* = 20 min, 40 min and 3.5 hr to the mean data points; Equations 3–13 were solved with depleting tear ranibizumab concentration, constant tear volume and in the absence of VEGF. **Bottom row**: a single injection is administered at *t* = 0 hr; simulations start at *t* = 20 min (= 1/3 hr) with *r*_Aq_(1/3) = 4.73×10^−2^ pmol ml^−1^ and *r*_Vit_(1/3) = 4.29 pmol ml^−1^, equal to the mean data points at those times; fitting was performed for *r*_Aq_ at *t* = 40 min, 1 hr and 3.5 hr to the mean data points; Equations 4–12 were solved in the absence of VEGF. Reasonable fits are achieved in all cases except for vitreal ranibizumab with topical treatment. Model fit 1: β_Tear Aq,*r*_ = 5.93 10^−7^ cm hr^−1^ (this value is also used for model fit 2 in the topical case) and β_Aq Vit,*r*_ = 0.929 cm hr^−1^; model fit 2: β_Aq Vit,*r*_ = 0.577 cm hr^−1^. All remaining parameters chosen as the default porcine values in Table 3.

Another fitting was explored, this time allowing fluid from the drop to flow off of the cornea (the tear volume loss case), and fitting for τ_loss_ in addition to β_Tear−Aq,*r*_ and β_Aq−Vit,*r*_. While a closer fit was obtained than for model fit 1, this would be expected when fitting a greater number of parameters, and the fit requires β_Aq−Vit,*r*_ ≈ 0 cm hr^−1^ which is unrealistic. Therefore, we consider model fit 1 to be preferable; the unrealistic fit indicating that the majority of the drop volume remains on the cornea during the experiment (a result consistent with our visual observations).

A further fitting was explored to the intravitreal injection without CPPs experimental data, for which a single injection is administered at *t* = 0 hr. Equations 4–12 were solved in the absence of VEGF. Simulations were initialised at *t* = 20 min (= 1/3 hr) with *r*_Aq_(1/3) = 4.73 × 10^−2^ pmol ml^−1^ and *r*_Vit_(1/3) = 4.29 pmol ml^−1^, equal to the mean data points at those times. We initialise at *t* = 20 min, rather than *t* = 0 hr, since the measured vitreal ranibizumab concentrations are lower than would be expected given the injected dose (see Experimental results). (These values are also more appropriate, since the measured values are more representative of the vitreal ranibizumab concentration near the aqueous-vitreous interface.) Fitting was performed for *r*_Aq_ at *t* = 40 min, 1 hr and 3.5 hr to the mean data points, the aqueous ranibizumab measurements being more informative than the vitreal measurements which remain roughly constant over time.

A reasonable fit was achieved in the aqueous and vitreous (*model fit 2*: β_Aq−Vit,*r*_ = 0.577 cm hr^−1^; see the bottom row of Fig. 4). The fitted value of β_Aq−Vit,*r*_ in model fit 2 is a factor of about 0.62 what it was in model fit 1. We plot simulation results using β_Aq−Vit,*r*_ values from both model fits for both topical and intravitreal treatment (using the model 1 fit value for β_Tear−Aq,*r*_ for model fit 2 in the topical case; Fig. 4). It can be seen that model fit 1 provides a good fit in both cases, and actually does a better job in fitting to the *t* = 3.5 hr data point in the aqueous for intravitreal treatment than model fit 2. Therefore, we consider model fit 1 to be preferable, taking this as our default for the *ex vivo* porcine eye.

Finally, we repeated each of the above fits, this time including VEGF. As anticipated, the effect on the fitted values was negligible given that VEGF concentrations are at least an order of magnitude lower than ranibizumab concentrations in each compartment.

### Human eyes

Having fitted our model to *ex vivo* porcine data in the previous section (Model fitting), we use our model, with appropriate modification of the equations (2–12) and parameters (see Model formulation, Table 3 and Appendix A: Parameter justification), to predict how ranibizumab will behave in the *in vivo* human eye and its effect upon VEGF levels.

### Model predictions

We begin by considering treatments involving a single mode of ranibizumab administration, either via topical drops (with CPP), drug-eluting contact lenses (with CPP) or intravitreal injections (without CPP; see Fig. 5); where CPP is not modelled explicitly, but assumed to be present for topical administration to allow ranibizumab to pass through the cornea.

**Figure 5:**
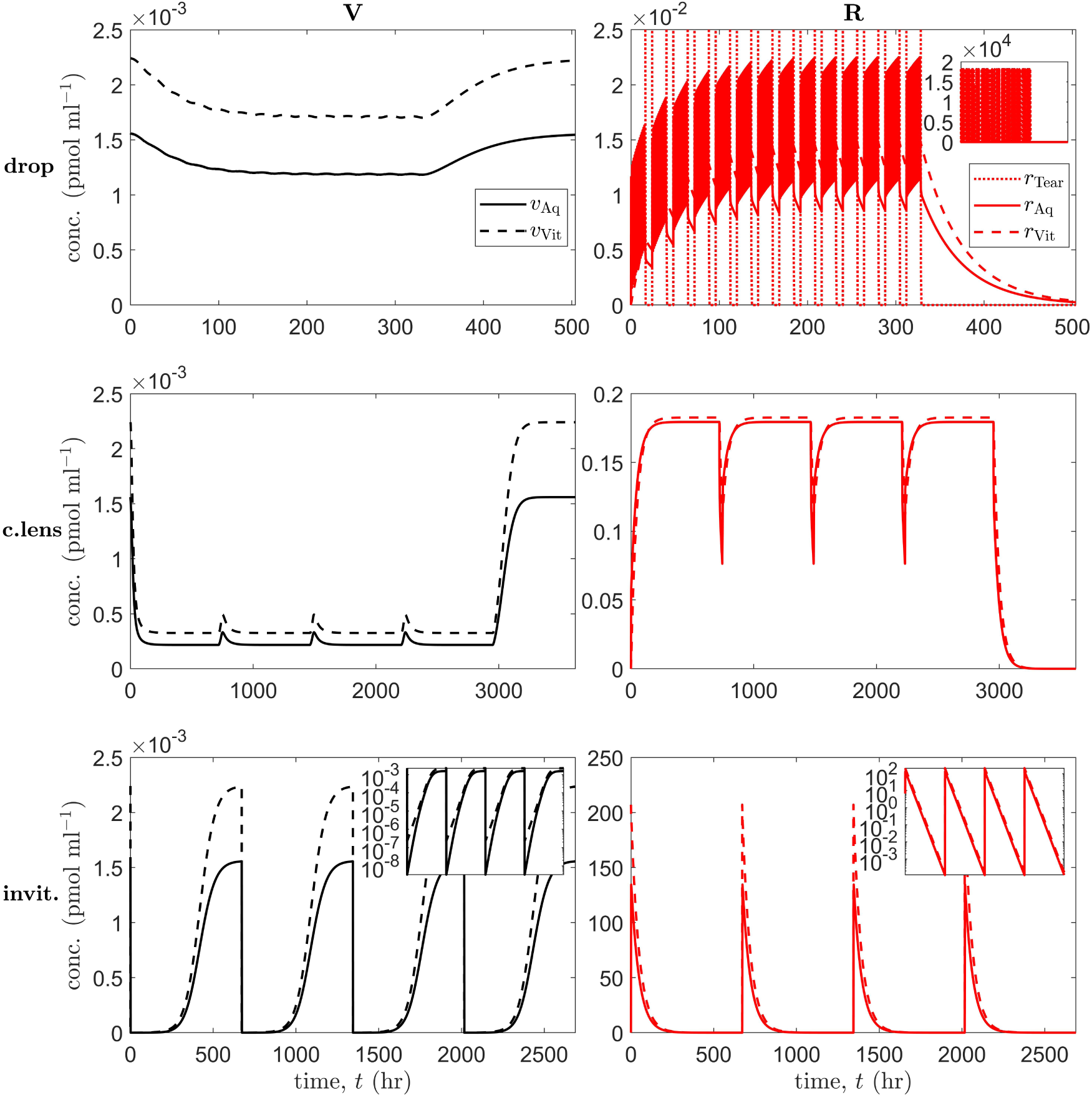
Simulation results for treatment of an *in vivo* human eye, using a single mode of administration. Panels show VEGF (V) concentrations (**left column**) and ranibizumab (R) concentrations (**right column**) in the tear film (Tear), aqueous (Aq) and vitreous (Vit), as appropriate (insets show results over the full range of concentrations (top-right), or with a logarithmic scale on the ordinate (bottom row)). **Top row** (topical drops): a single drop is applied at the start of hours 1–16 every day for the first 2 weeks; week 3 untreated. **Middle row** (drug-eluting contact lens): a series of 4 lenses are worn for 30 days at a time, starting on day 1, with a 1 day break between lenses; final 4 weeks untreated. **Bottom row** (intravitreal injections): administered at the start of weeks 1, 5, 9 and 13; simulation runs to 16 weeks. Drops suppress aqueous and vitreal VEGF levels to a fairly constant value during the period of administration; contact lenses suppress VEGF levels more strongly, with small transient increases in VEGF levels between lenses; injections reduce VEGF to by far the lowest levels, though VEGF returns to untreated values between injections. Equations 2–12 were solved using the default human parameters in Table 3.

First, in the topical drop case, we consider the scenario where a single drop is applied at the start of hours 1–16 every day for the first 2 weeks, with week 3 left untreated (Fig. 5 (top row)). This assumes the patient is awake 16 hours a day and applies the treatment once each waking hour. Though 4 drops a day would be a more realistic maximum frequency, ^71^ the aim here is to explore the maximum theoretical effect from topical drops. Aqueous and vitreal VEGF levels are suppressed to fairly constant values during the period of administration (1.18 × 10^−3^ and 1.70 × 10^−3^ pmol ml^−1^ respectively), reaching suppressed levels after about 100 hours of treatment, and returning to untreated values (1.56 × 10^−3^ and 2.24 × 10^−3^ pmol ml^−1^ respectively) after about a week post treatment termination, while aqueous and vitreal ranibizumab concentrations oscillate over ranges of (8.57 × 10^−3^, 2.25 × 10^−2^) and (1.25 × 10^−2^, 1.48 × 10^−2^) pmol ml^−1^ respectively, once a regular periodic pattern has been established.

Second, in the drug-eluting contact lens case, we consider the scenario where a series of 4 lenses are worn for 30 days at a time (a realistic maximum wear time for continuous wear contact lenses; see, for example, www.specsavers.co.uk/contact-lenses and Lin et al. ^72^), starting on day 1, with a 1 day break between lenses, with the final 4 weeks left untreated (Fig. 5 (middle row)). VEGF levels were suppressed more strongly than with drops (with periodic minima of 2.17 × 10^−4^ and 3.25 × 10^−4^ pmol ml^−1^ in the aqueous and vitreous respectively), with small transient increases in VEGF levels between lenses (with periodic maxima of 3.32 × 10^−4^ and 4.93 × 10^−4^ pmol ml^−1^ in the aqueous and vitreous respectively). The initial minimum is achieved after about 150 hours of treatment, and VEGF levels return to untreated values after about 2 weeks post treatment termination. Aqueous and vitreal ranibizumab concentrations oscillate over ranges of (7.64 × 10^−2^, 0.180) and (0.111, 0.183) pmol ml^−1^ respectively once a regular periodic pattern has been established.

Third, in the intravitreal injection case, we consider the scenario where injections are administered every 4 weeks (i.e. monthly, this being the highest frequency at which anti-VEGF injections are generally administered, ^73,74^ the aim being to show the maximum effect), at the start of weeks 1, 5, 9 and 13, with the simulation allowed to run to the end of week 16 (Fig. 5 (bottom row)). Injections reduce VEGF to by far the lowest levels (3.27 × 10^−9^ and 2.21 × 10^−7^ pmol ml^−1^ in the aqueous and vitreous respectively), though VEGF returns to untreated values between injections. VEGF levels reach their minimum about 1.5 hours after injection in the aqueous and 12 minutes after injection in the vitreous, and return to untreated values after about 4 weeks post treatment termination. Aqueous and vitreal ranibizumab concentrations oscillate over ranges of (1.13 × 10^−4^, 135) and (1.64 × 10^−4^, 207) pmol ml^−1^ respectively.

The above results demonstrate that topical treatments have the advantage of maintaining VEGF at suppressed levels, while intravitreal treatments reduce VEGF levels far more significantly, with the disadvantage that VEGF levels return to untreated values between treatments. Therefore, we considered two further cases: dual drop/intravitreal administration and dual contact lens/intravitreal administration (see Fig. 6).

**Figure 6:**
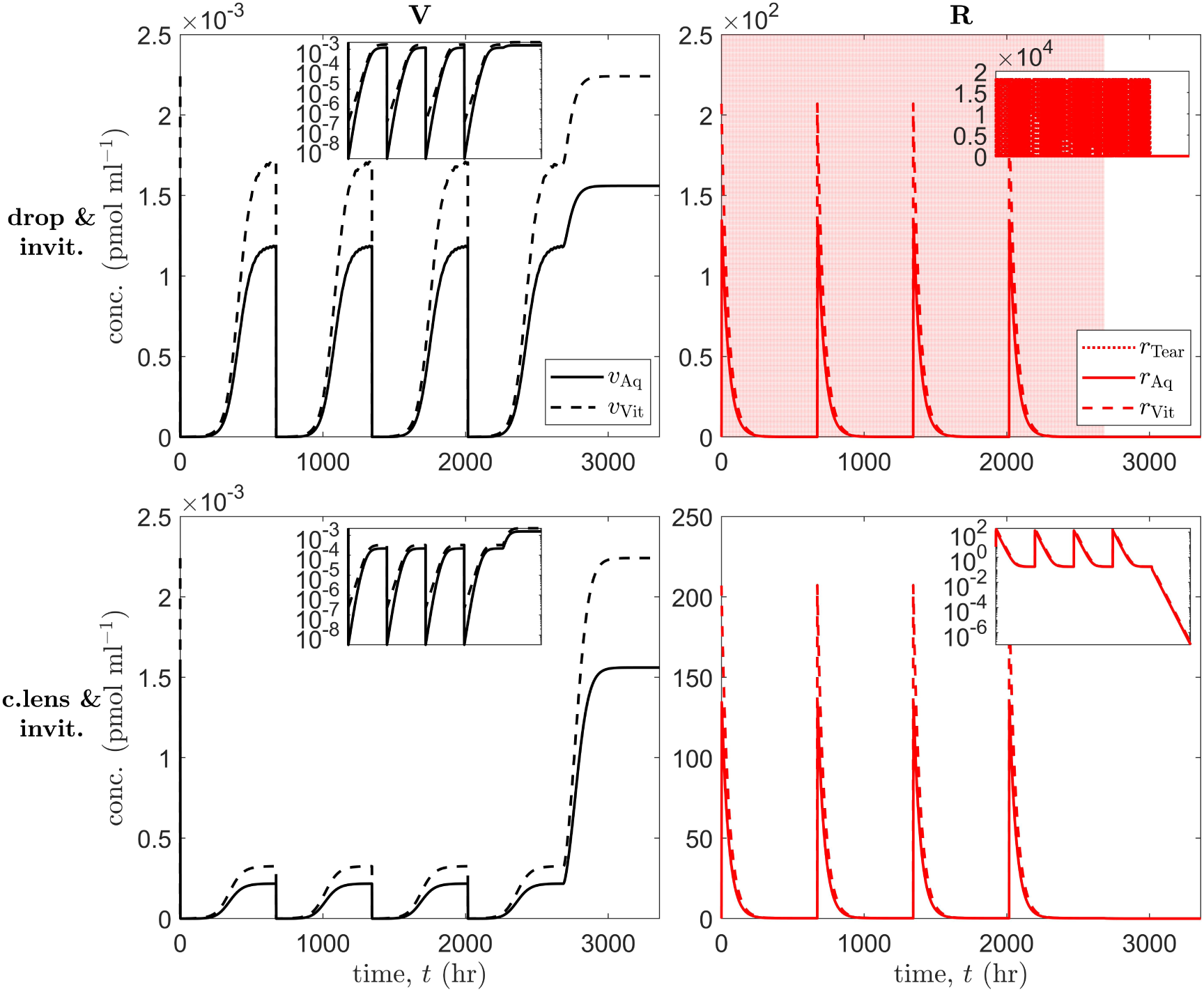
Simulation results for treatment of an *in vivo* human eye, using multiple modes of administration. Panels show VEGF (V) concentrations (**left column**) and ranibizumab (R) concentrations (**right column**) in the tear film (Tear), aqueous (Aq) and vitreous (Vit), as appropriate (insets show results over the full range of concentrations (top-right), or with a logarithmic scale on the ordinate (top-left and bottom row)). **Top row** (topical drops and intravitreal injections): a single drop is applied at the start of hours 1–16 every day for the first 16 weeks, while injections are administered at the start of weeks 1, 5, 9 and 13; simulation runs to 20 weeks, the final 4 untreated. **Bottom row** (drug-eluting contact lens and intravitreal injections): a series of 4 lenses are worn for 27 days at a time, starting on day 2, with a 1 day break between lenses, while injections are administered at the start of weeks 1, 5, 9 and 13 (on the days without contact lenses); simulation runs to 20 weeks, the final 4 untreated. Both drops and drug-eluting lenses suppress aqueous and vitreal VEGF levels between injections, preventing them from returning to untreated levels as in Fig. 5 (bottom row). Equations 2–12 were solved using the default human parameters in Table 3.

First, for dual drop/intravitreal administration, we consider the scenario where a single drop is applied at the start of hours 1–16 every day for the first 16 weeks, while injections are administered at the start of weeks 1, 5, 9 and 13 (Fig. 6 (top row)). The simulation runs to 20 weeks, the final 4 of which are untreated. VEGF levels remain damped between injections, aqueous and vitreal concentrations oscillating over ranges of (3.28 × 10^−9^, 1.19 × 10^−3^) and (2.21 × 10^−7^, 1.71 × 10^−3^) pmol ml^−1^ respectively, while aqueous and vitreal ranibizumab concentrations oscillate over ranges of (8.70 × 10^−3^, 135) and (1.26 × 10^−2^, 207) pmol ml^−1^ respectively.

Finally, for dual contact lens/intravitreal administration, we consider the scenario where a series of 4 lenses are worn for 27 days at a time, starting on day 2, with a 1 day break between lenses, while injections are administered at the start of weeks 1, 5, 9 and 13 (on the days without contact lenses; Fig. 6 (bottom row)). The simulation runs to 20 weeks, the final 4 of which are untreated. VEGF levels are damped more strongly than with drops between injections, aqueous and vitreal concentrations oscillating over ranges of (3.29 × 10^−9^, 2.17 × 10^−4^) and (2.20 × 10^−7^, 3.25 × 10^−4^) pmol ml^−1^ respectively, while aqueous and vitreal ranibizumab concentrations oscillate over ranges of (0.180, 135) and (0.183, 207) pmol ml^−1^ respectively.

### Sensitivity analysis

A local sensitivity analysis was performed to determine the effect of varying each parameter upon key model outputs. Parameters were varied individually, across 101 values uniformly distributed over the biologically realistic ranges given in Table 5, while keeping all remaining parameters fixed at their default values in Table 5. For each parameter set, Equations 2–12 were solved for *t* ∈ [0, 12] weeks and the outputs from the interval *t* ∈ [9, 12] weeks — within which the solution has firmly settled to either an unchanging steady-state or a regular periodic pattern — used to calculate the maximum, mean and minimum vitreal values of V and R_Tot_ (= R + VR + 2RVR, the total ranibizumab concentration; see Supplementary Figs. S1–S6). We chose vitreal values as our key outputs since these are the most relevant in preventing choroidal neovascularisation, and calculate the max/mean/min since it is important to minimise all three quantities for VEGF and to maximise all three quantities for R_Tot_. We use R_Tot_ rather than R since the total quantity of ranibizumab is of greater significance than the free quantity — sensitivity analysis was also performed on R and the results were almost identical to those for R_Tot_.

**Table 5:** Parameter default values and value ranges for sensitivity analysis of the *in vivo* human model.

| Parameter | Default value | Range of values |
| --- | --- | --- |
| $\tau_{\text{loss}}$ | $5.00 \times 10^{-2}$ hr | $[1.67, 8.33] \times 10^{-2}$ hr |
| $V_{\text{Drop}}$ | $4.50 \times 10^{-2}$ ml | $[4.05, 4.95] \times 10^{-2}$ ml |
| $V_{\text{TearNorm}}$ | $6.35 \times 10^{-3}$ ml | $[0.340, 1.07] \times 10^{-2}$ ml |
| $V_{\text{TearRes}}$ | $3.00 \times 10^{-2}$ ml | $[2.70, 3.30] \times 10^{-2}$ ml |
| $V_{\text{Aq}}$ | 0.160 ml | $[0.121, 0.286]$ ml |
| $V_{\text{Vit}}$ | 4.50 ml | $[3.50, 5.40]$ ml |
| $A_{\text{Tear-Aq}}$ | 1.30 cm<sup>2</sup> | $[1.04, 1.56]$ cm<sup>2</sup> |
| $A_{\text{Aq-Vit}}$ | 0.349 cm<sup>2</sup> | $[0.111, 0.521]$ cm<sup>2</sup> |
| $A_{\text{Vit-Ret}}$ | 10.9 cm<sup>2</sup> | $[9.81, 12.0]$ cm<sup>2</sup> |
| $k^+$ | $0.576 \text{ pmol}^{-1} \text{ ml hr}^{-1}$ | $[0.205, 4.21] \text{ pmol}^{-1} \text{ ml hr}^{-1}$ |
| $k^-$ | $2.63 \times 10^{-2} \text{ hr}^{-1}$ | $[1.40, 3.60] \times 10^{-2} \text{ hr}^{-1}$ |
| $\delta_{i,j}^*$ | 0 hr<sup>-1</sup> | $[0, 0.770] \text{ hr}^{-1}$ |
| $\beta_{\text{Tear-Aq},r}$ | $1.10 \times 10^{-6} \text{ cm hr}^{-1}$ | $[0.990, 1.21] \times 10^{-6} \text{ cm hr}^{-1}$ |
| $\beta_{\text{Aq-Vit},v}$ | 0.985 cm hr<sup>-1</sup> | $[0.887, 1.08] \text{ cm hr}^{-1}$ |
| $\beta_{\text{Aq-Vit},r}$ | 0.929 cm hr<sup>-1</sup> | $[0.836, 1.02] \text{ cm hr}^{-1}$ |
| $\beta_{\text{Aq-Vit},u}$ | 0.755 cm hr<sup>-1</sup> | $[0.680, 0.831] \text{ cm hr}^{-1}$ |
| $\beta_{\text{Aq-Vit},w}$ | 0.653 cm hr<sup>-1</sup> | $[0.588, 0.718] \text{ cm hr}^{-1}$ |
| $\beta_{\text{Vit-Ret},r}$ | $6.80 \times 10^{-4} \text{ cm hr}^{-1}$ | $[6.12, 7.48] \times 10^{-4} \text{ cm hr}^{-1}$ |
| $\beta_{\text{Vit-Ret},u}$ | $6.44 \times 10^{-4} \text{ cm hr}^{-1}$ | $[5.80, 7.08] \times 10^{-4} \text{ cm hr}^{-1}$ |
| $\beta_{\text{Vit-Ret},w}$ | $6.23 \times 10^{-4} \text{ cm hr}^{-1}$ | $[5.61, 6.85] \times 10^{-4} \text{ cm hr}^{-1}$ |
| $\phi_{\text{Vit},v}$ | $2.34 \times 10^{-4} \text{ pmol hr}^{-1}$ | $[1.02, 4.46] \times 10^{-4} \text{ pmol hr}^{-1}$ |
| $\psi_{\text{Tear}}$ | $7.20 \times 10^{-2} \text{ ml hr}^{-1}$ | $[0.300, 1.32] \times 10^{-1} \text{ ml hr}^{-1}$ |
| $\psi_{\text{Aq}}$ | 0.150 ml hr<sup>-1</sup> | $[0.660, 2.52] \times 10^{-1} \text{ ml hr}^{-1}$ |
| $r_{\text{Dose}}$ | $2.07 \times 10^4 \text{ pmol ml}^{-1}$ | $[1.86, 2.28] \times 10^4 \text{ pmol ml}^{-1}$ |

See Appendix A: Parameter justification for explanation of parameter values.
\*: for $i \in \{\text{Aq}, \text{Vit}\}$ and $j \in \{v, r, u, w\}$ .

For each key output and parameter, we calculate the *sensitivity factor*, defined as the maximum value obtained by the output over the 101 values across which the parameter was varied, divided by the minimum value over that range. Thus, for a given treatment scenario, each parameter has 6 sensitivity factors associated with it, one for each of the key model outputs (max/mean/min of V/R_Tot_). We consider a sensitivity factor to be significant if it exceeds 1.5, such that the key model output varies by over 50% of its minimum value within the range of values considered for that parameter.

Each mode of administration was considered in isolation: topical drops, applied on the hour, every hour (Fig. 7 (top row)); a drug-eluting contact lens, worn continuously (Fig. 7 (middle row)); and intravitreal injections, administered at the start of weeks 1, 5, and 9 (Fig. 7 (bottom row)). (While, in practice, drops would not be administered every hour of the day, and a continuous wear contact lens could be worn for a maximum of 30 days, this is not important for the purposes of our sensitivity analysis.) For topical drops, the model demonstrates sensitivity to: (*V*_TearNorm_, *A*_Aq−Vit_, *k*^+^, δ_Aq,*r*_, δ_Vit,*v*_, δ_Vit,*r*_, φ_Vit,*v*_, ψ_Tear_, ψ_Aq_); for a drug-eluting contact lens, the model demonstrates sensitivity to: (*A*_Tear−Aq_, *A*_Aq−Vit_, *k*^+^, δ_Aq,*r*_, δ_Vit,*v*_, δ_Vit,*r*_, φ_Vit,*v*_, ψ_Aq_); and for intravitreal injections, the model demonstrates sensitivity to: (*V*_Vit_, *A*_Aq−Vit_, *k*^+^, δ_Aq,*r*_, δ_Vit,*v*_, δ_Vit,*r*_, δ_Vit,*u*_, β_Aq−Vit,*r*_, φ_Vit,*v*_, ψ_Aq_). See Supplementary Figs. S1–S6, for detailed plots showing the variation of each key output with each parameter under each mode of administration.

**Figure 7:**
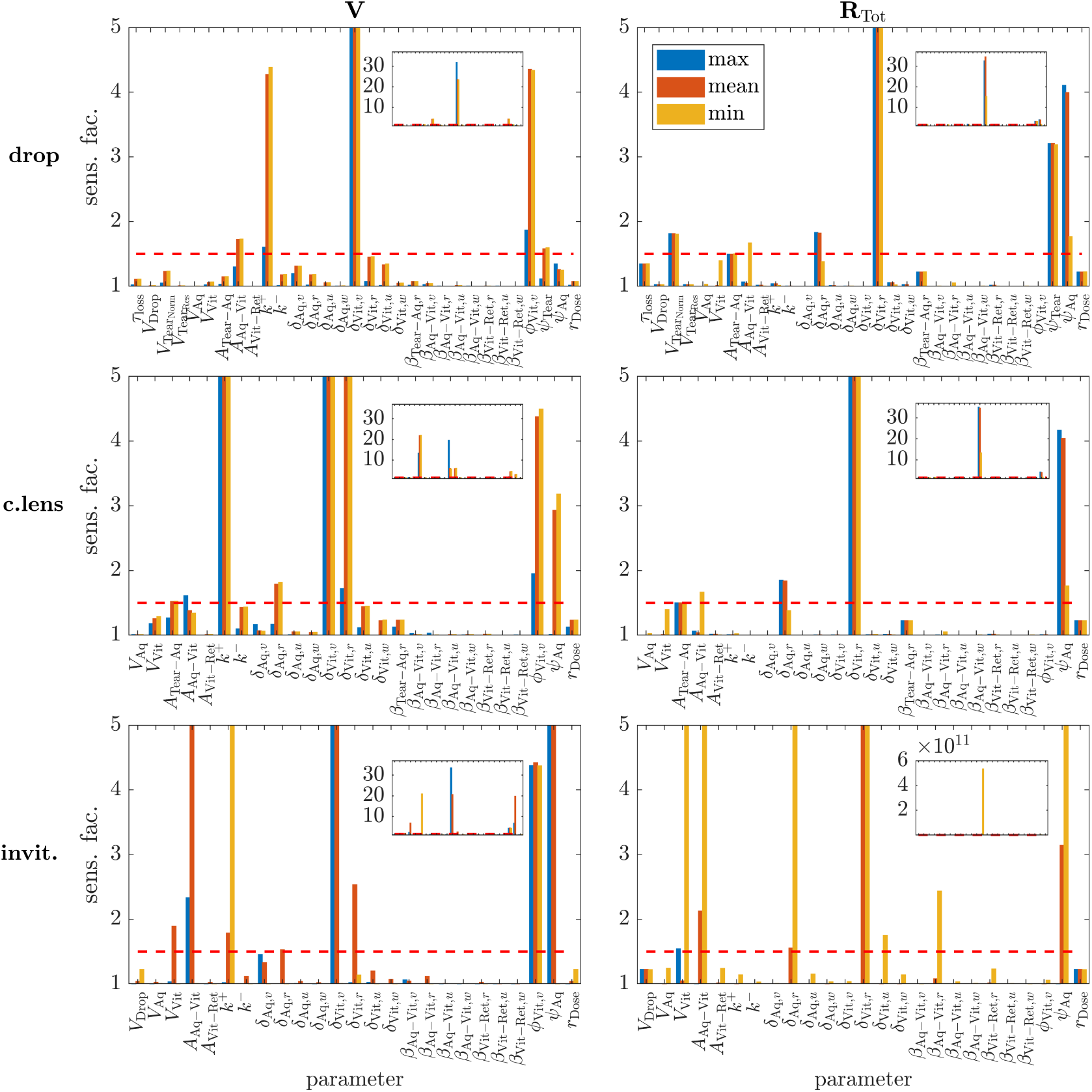
Local sensitivity analysis. Panels show sensitivity of the maximum/mean/minimum vitreal VEGF (V) concentration (**left column**) and total vitreal ranibizumab (R_Tot_ = R + VR + 2RVR) concentration (**right column**) to variation in model parameters over biologically realistic ranges (insets show full range of sensitivity values). Equations 2–12 were solved for *t* ∈ [0, 12] weeks. **Top row**: topical drops, applied on the hour, every hour. **Middle row**: drug-eluting contact lens, worn continuously. **Bottom row**: intravitreal injections, administered at the start of weeks 1, 5, and 9. Parameters were varied individually, across 101 values uniformly distributed over the ranges given in Table 5, the remaining parameters being held at their default values given in Table 5. For each parameter set, the maximum/mean/minimum vitreal values of V and R_Tot_ were calculated over model outputs from the interval *t* ∈ [9, 12] weeks (see Supplementary Figs. S1–S6). Each sensitivity factor was then calculated as the maximum value obtained by the maximum/mean/minimum value of V or R_Tot_ over the 101 values across which the parameter was varied, divided by the corresponding minimum value over that range. The dashed red horizontal line demarcates the sensitivity threshold (=1.5), above which sensitivity is considered significant. The model demonstrates sensitivity to the following parameters — drops: (*V*_TearNorm_, *A*_Aq−Vit_, *k*^+^, δ_Aq,*r*_, δ_Vit,*v*_, δ_Vit,*r*_, φ_Vit,*v*_, ψ_Tear_, ψ_Aq_); contact lens: (*A*_Tear−Aq_, *A*_Aq−Vit_, *k*_+_, δ_Aq,_*_r_*, δ_Vit,_*_v_*, δ_Vit,_*_r_*, φ_Vit,_*_v_*, ψ_Aq_), injections: (*V*_Vit_, *A*_Aq−Vit_, *k*_+_, δ_Aq,_*_r_*, δ_Vit,_*_v_*, δ_Vit,_*_r_*, δ_Vit,_*_u_*, β_Aq−Vit,_*_r_*, φ_Vit,_*_v_*, ψ_Aq_).

## Discussion

Topical application of anti-VEGF drugs provides a promising alternative or addition to intravitreal injections, which are expensive, uncomfortable and risk infection. In this study, we took a combined experimental-modelling approach to investigate the potential for topical ranibizumab administration in the treatment of wet-AMD.

Our experiments in *ex vivo* porcine eyes revealed that our (poly-arginine based) CPP allows topical ranibizumab to penetrate the cornea (in agreement with de Cogan et al. ^9^), though it reduces ranibizumab’s efficacy in neutralising VEGF for intravitreal treatment. This latter effect is likely due to CPP reducing ranibizumab’s ability to bind to VEGF in the time interval, post administration, during which CPPs unbind from ranibizumab. As would be expected, intravitreal ranibizumab treatments result in higher vitreal ranibizumab levels, and more rapid and complete reduction of vitreal VEGF than topical treatment.

Mathematical modelling allowed us both to gain a deeper insight into our experimental results, and to extrapolate to the living human eye. Fitting our model to the experimental data, we determined the permeabilities of the porcine cornea and suspensory ligaments (zonule) to ranibizumab, VEGF and their compounds (RV and RVR), extrapolating these values for the human. Further, our model fits suggest that a substantive proportion of the drop volume remains on the eye during experiments, consistent with visual observations.

In the *in vivo* human eye, we used our mathematical model to simulate ranibizumab and VEGF levels (and those of their compounds) in the tear film, aqueous and vitreous, under a number of different treatment regimens. We note that, given our model does not contain retinal or choroidal compartments, the best measure of each treatment’s efficacy is its effect on vitreal VEGF levels. First, we considered cases where treatment is applied via a single mode of administration: drops, drug-eluting contact lenses or intravitreal injections. We found that topical therapies (drops and contact lenses) maintain suppressed vitreal VEGF levels (and maintain vitreal ranibizumab levels), drug-eluting contact lenses having a stronger suppressive effect given that they maintain a constant supply of ranibizumab to the tear film, whereas ranibizumab delivered via drops is quickly diluted and removed from the tear. By comparison, intravitreal injections suppress vitreal VEGF levels more strongly, as would be expected given that a large ranibizumab dose is delivered directly to the vitreous; however, the effect wears off between treatments, vitreal VEGF levels returning to untreated values between treatments in a 4-week regimen. This is on the lower end of the range of VEGF suppression periods (26–69 days) measured in humans following an intravitreal ranibizumab injection. ^75^

If topical treatment were able to maintain vitreal (and, hence, retinal/choroidal) VEGF levels at sufficiently low concentrations, then topical therapy could replace intravitreal therapy. Even if this is not possible, topical modes of administration could potentially be used in combination with intravitreal treatments, increasing the overall efficacy of the treatment and reducing the frequency with which intravitreal injections need to be administered. To this end, we simulated dual topical and intravitreal administration, combining either drops and injections, or drug-eluting contact lenses and injections. Our simulations suggest that dual therapies would provide the best of both worlds, with injections providing highly suppressed vitreal VEGF levels, and topical treatment preventing VEGF levels from returning to untreated values between injections (such that VEGF levels do no exceed the suppressed levels achieved by applying topical treatment alone).

Local sensitivity analysis revealed the parameters to which the vitreal ranibizumab and VEGF concentrations are most sensitive. Some of these parameters, namely *V*_TearNorm_, *V*_Vit_, *A*_Tear−Aq_, *A*_Aq−Vit_, δ_Vit,*v*_, δ_Vit,*u*_, φ_Vit,*v*_ and ψ_Aq_, may vary between patients, though they are not factors that we could realistically alter, being fundamental to the eye’s geometry, biochemistry or biomechanics. However, other parameters, namely *k*^+^, δ_Aq,*r*_, δ_Vit,*r*_, β_Aq−Vit,*r*_ and ψ_Tear_, could potentially be modified by redesigning the drug or mode of application. For example, *k*^+^ could perhaps be increased, and β_Aq−Vit,*r*_ (for intravitreal delivery), δ_Aq,*r*_ and δ_Vit,*r*_ decreased, by altering the drug’s molecular structure, while ψ_Tear_ could be decreased by increasing ranibizumab’s residence time in the tear film, e.g. using a molecular anchor such as collagen binding domains ^76^ or wheat germ agglutinin, ^77^ or viscosity increasing polymers such as polyacrylates and polyvinyl alcohols. ^39^

In future studies, we will extend this work on both experimental and mathematical fronts. Future experimental work could include measuring retinal ranibizumab and VEFG concentrations in addition to those in the aqueous and vitreous, measuring CPP concentrations in each compartment, making more frequent measurements over a larger time span with more repeats, and measurements using *in vivo* animal eyes in which key elements such as clearance mechanisms and VEGF production are active. Preferably, this would be done in eyes of a similar size to human eyes such as porcine eyes, though rat eyes would also be informative. It would also be interesting to repeat these measurements with drug-eluting contact lenses and to explore ways to further improve ranibizumab’s corneal penetration and tear residence time, and their effect on treatment efficacy. Future mathematical modelling will include extending our ODE model to include a retinal compartment (in a similar way to Hutton-Smith et al. ^68^), and developing a spatially-resolved partial differential equation model accounting for both the advective and diffusive transport of ranibizumab, VEGF and their compounds.

The combined experimental-modelling approach taken in this paper has provided deeper insights than would be possible by employing either approach in isolation. Our experimental results and modelling predictions suggest that topical application may be a promising means of ranibizumab administration, either in isolation, or in combination with intravitreal injections; reducing the frequency of injections and improving overall treatment efficacy. While formulated with ranibizumab in mind, our model could easily be adapted to simulate topical and intravitreal administration of other anti-VEFG drugs, and of ocular drugs in general, including new and upcoming treatments for non-neovascular AMD such as pegcetacoplan and avacincaptad pegol. ^23–25^ Using our mathematical model and variations upon it, we can accelerate the development of new drugs, administration techniques and regimens, testing a range of scenarios quickly and at low cost, as well as reducing, replacing and refining animal experiments.

## Appendix A: Parameter justification

• **Time for tear volume to return to normal (human)** / time for eye drop to leave cornea (porcine), τ_loss_

**–** *In vivo* human eye: estimates of the time taken for the tear volume to return to normal following the application of a drop range between 1–5 min = 1.67–8.33 × 10^−2^ hr. ^37–41^ We assume that τ_loss_ could lie anywhere within this range, taking the mean value of 3 min = 5 × 10^−2^ hr as our default.

**–** *Ex vivo* porcine eye: it is difficult to characterise the time taken for a drop to flow off of the cornea; however, observations during our experiments suggest that the majority of the drop volume remains on the cornea during the experiment (the constant volume scenario). When considering the depleting volume scenario, we assume that the timescale is similar to that for the human tear volume to return to normal, given above. Therefore, we take a value of 3 min = 5 × 10^−2^ hr as our default, with a possible range of 1.67–8.33 × 10^−2^ hr.

• **Volume of eye drop (human and porcine),** *V*_Drop_: the volume of the eye drop used in our experiments is 4.5 × 10^−2^ ml and we assume the same volume for human treatment. For the sensitivity analysis, we assume that *V*_Drop_ could vary over a range 10% above and below its default value, giving the range 4.05–4.95 × 10^−2^ ml.

• **Volume of normal tear film (human),** *V*_TearNorm_: the normal volume of the human tear film has been measured to range between 3.4–10.7 × 10^−3^ ml. ^42,43^ We assume that *V*_TearNorm_ could lie anywhere within this range, taking the mean of the mean values stated in Mishima et al. ^42^ and Scherz et al. ^43^, that is 6.35 × 10^−3^ ml, as our default value (see also 37, 38, 40, 44, 45, 78–80).

• **Volume of tear reservoir (human),** *V*_TearRes_: the human tear reservoir volume has been measured to be 3 × 10^−2^ ml. ^38,40,44,45^ For the sensitivity analysis, we assume that *V*_TearRes_ could vary over a range 10% above and below its default value, giving the range 2.70–3.30 × 10^−2^ ml.

_•_ **Volume of aqueous, *V*_Aq_**

**–** *In vivo* human eye: the volume of the aqueous in the healthy human eye decreases with age, with values of 0.247 ± 0.039 ml in people aged 20–30 yrs and values of 0.160 ± 0.039 ml in people aged 60 yrs and older. ^46^ We take a value of 0.160 ml as our default, since patients being treated for wet AMD will typically be 60 yrs and older, and assume that *V*_Aq_ could lie anywhere in the range 0.121–0.286 ml (where, 0.121 = 0.160 − 0.039 and 0.286 = 0.247 + 0.039; see also 17, 81).

**–** *Ex vivo* porcine eye: the aqueous volume of the pig eye has been measured to be 0.31 ml. ^47,48^

**_•_ Volume of vitreous, *V*_Vit_**

**–** *In vivo* human eye: the volume of the vitreous has been measured to lie in the range 3.5–5.4 ml, ^47^ with quoted volumes of 4 ml ^52^ and 4.5 ml ^17,49,50^ being common. We assume that *V*_Vit_ could lie anywhere within the range 3.5–5.4 ml, taking a value of 4.5 ml as our default since it lies in the middle of this range and is a frequently quoted value.

**–** *Ex vivo* porcine eye: the vitreal volume of the pig eye has been measured to lie in the range 3.0–3.2 ml. ^47,48,51,52^ We assume that *V*_Vit_ could lie anywhere within this range, taking the middle value of 3.1 ml as our default.

**_•_ Area of the tear-aqueous interface, *A*_Tear−Aq_**

**–** *In vivo* human eye: Rüfer et al. ^54^ measured the corneal diameter of human eyes to range across 1.07–1.26 cm, with an average diameter of 1.17 cm, while Bron et al. ^53^ quote a horizontal diameter of 1.175 cm and a vertical diameter of 1.06cm. Bron et al. also give the corneal anterior radius of curvature as 0.78 cm. Approximating the cornea as a flat disc, Rüfer et al.’s measurements correspond to corneal areas in the range 0.899–1.24 cm^2^ with a mean value of 1.08 cm^2^, while approximating the cornea as a flat ellipse, Bron et al.’s measurements correspond to an area of 0.978 cm^2^. If we instead use a surface integral to calculate the corneal anterior area, using the corneal anterior radius of curvature, then Rüfer et al.’s measurements correspond to areas ranging across 1.04–1.56 cm^2^ with a mean value of 1.30 cm^2^, while Bron et al.’s measurements correspond to an area of 1.16 cm^2^. The corneal area quoted by Bron et al. is 1.06 cm^2^. This is smaller than our calculated area based on Bron et al.’s diameters, but within the range of areas calculated from Rüfer et al.’s diameters. We assume that corneal area may range across 1.04–1.56 cm^2^ and take 1.30 cm^2^ as our default value.

**–** *Ex vivo* porcine eye: the corneal surface area of ‘large’ porcine eyes has been measured to be 1.40 ± 0.19 cm^2^. ^55^ We scale this value for eyes of the dimensions used in our experiments as follows. The mean scleral surface area of the ‘large’ porcine eyes noted above was measured to be 11.92 cm^2, 55^ giving a total mean surface area of 13.32 cm^2^ (assuming that the limbal surface area is included in the scleral surface area). Thus, the proportion of the eye surface area taken up by the cornea is 0.105 (this proportion would appear to be scale-invariant, given that it is maintained for their measurements of ‘small-’ and ‘medium’-sized pig eyes –– these being 0.123 and 0.106 respectively). We measured the length and width of each of the eyes used in our experiments (providing similar values to those given in previous studies ^47,62^). Taking the average of these two diameters for each eye and approximating the eye’s geometry as a sphere, we calculated each eye’s surface area. This gave a mean total surface area across all eyes of 14.40 cm^2^, with a range of 12.33–16.74 cm^2^. Assuming that the proportion of the eye’s surface area taken up by the cornea is the same as in Olsen et al. ^55^, we take the default corneal surface area to be 1.51 cm^2^, with a possible range of 1.30-1.76 cm^2^. This is also within the range of values which can be extrapolated from measurements in Heichel et al. ^82^ and Sanchez et al. ^61^.

**_•_ Area of the aqueous-vitreous interface, *A*_Aq−Vit_**

**–** *In vivo* human eye: the aqueous-vitreous interface consists of the suspensory ligaments which span an annular region between the ciliary body and the lens (holding it in place). We calculate its area as follows. Manns et al. ^56^ measured the lens diameter and inner ciliary body diameter in 3 groups of individuals: a young group (8–19 years), an older group of prepresbyopic donors (20-–70 years) and an older group of presbyopic donors (20-–70 years). The mean lens diameter across all three groups was 0.90 cm and ranged between 0.84–0.97 cm, while the mean inner ciliary body diameter across all three groups was 1.12 cm and ranged between 1.04–1.17 cm. This corresponds to lens areas ranging between 0.554–0.739 cm^2^, with a mean of 0.636 cm^2^, and areas spanned by the inner ciliary body ranging between 0.849–1.08 cm^2^, with a mean of 0.985 cm^2^. Thus, the mean aqueous-vitreous interface area ranges between 0.111–0.521 cm^2^ (taking this as our sensitivity analysis range), with a mean (default) value of 0.349 cm^2^ (which is also the mean area of the older prepresbyopic group).

**–** *Ex vivo* porcine eye: in the absence of an inner ciliary body diameter measurement, we calculate its area as follows. The scleral outer radius is 1.112 cm ^62^ (calculated to match the area value in Olsen et al. ^55^). The combined width of the retina and choroid varies across the eye, with a range of 0.02–0.086 cm ^53,61,62^, while the scleral width is 0.043–0.089 cm. ^55,62^ Subtracting the mean values of the retina-choroid and scleral widths from the scleral outer radius gives us the radius of the vitreous as 0.993 cm. (We note that this is consistent with the vitreal volume stated above, which, assuming a spherical vitreous, corresponds to a radius of 0.905 cm.) The length of the eye (in the transverse/sagittal planes) is 2.29 cm, ^62^ the corneal width is 0.098 cm at its apex., ^57,62^ and the anterior segment depth (that is, the lens equator position, posterior to the posterior surface of the cornea) is 0.247 cm. ^58,60,62^ Subtracting the scleral outer radius, the apex corneal width and the anterior segment depth from the length of the eye gives us the distance between the centre of the vitreous and the lens equator as 0.833 cm. Using Pythagoras’ theorem, this gives the radius of the aqueous-vitreous interface including the lens as 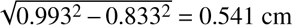. Therefore, the area of the aqueous-vitreous interface including the lens is 0.918 cm^2^. The lens has (unstretched) diameter of 0.886 cm ^59^ and hence an area of 0.617 cm^2^. Therefore, the area of the aqueous-vitreous interface, not including the lens, is 0.918 − 0.617 = 0.301 cm^2^.

• **Area of the vitreo-retinal interface (human),** *A*_Vit−Ret_: this has been calculated to be 10.9 cm^2^ ^63^ (our default value) and is similar to the value given for the choroidal surface area of 11.8 cm^2^ by Bron et al. ^53^ For the sensitivity analysis, we assume that *A*_Vit−Ret_ could vary over a range 10% above and below its default value, giving the range 9.81–12.0 cm^2^.

• **Binding rate of R to VR (to form RVR — human and porcine),** *k*^+^: Papadopoulos et al. ^64^ have measured a binding rate of ranibizumab to VEGF of 1.60 × 10^5^ M^−1^ s^−1^, converting to units consistent with our model, this is 0.576 pmol^−1^ ml hr^−1^ (our default value). This is within the range of values measured by Yang et al. ^83^, 0.205–4.21 pmol^−1^ ml hr^−1^ (following conversion, which we take as our range), and similar to the default value used by Hutton-Smith et al. ^17^

• **Unbinding rate of VR (to form V** + **R — human and porcine),** *k*^−^: Papadopoulos et al. ^64^ have measured an unbinding rate of ranibizumab from VEGF of 7.30 × 10^−6^ s^−1^, converting to units consistent with our model, this is 2.63×10^−2^ hr^−1^ (our default value). This is within the range of values measured by Yang et al. ^83^, 1.40–3.60×10^−2^ hr^−1^ (following conversion, which we take as our range), and similar to the default value used by Hutton-Smith et al. ^17^ The default values of *k*^+^ and *k*^−^ give us an equilibrium dissociation constant, 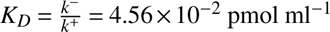. This is the same value as is used by Hutton-Smith et al. ^17^ and within the range of values resulting from Yang et al.’s ^83^ measurements.

• **Rate of molecular degradation,** δ*_i_*, *_j_* **(for** *i* ∈ {Aq, Vit} **and** *j* ∈ {*v*, *r*, *u*, *w*} **— human and porcine):** as a simplifying assumption, we neglect molecular degradation by default, assuming that it is negligible compared to the speed of the other processes captured in our model in the *in vivo* human eye and absent in the *ex vivo* porcine eye (indeed, molecular degradation is not typically included in models of anti-VEFG treatments; see, for example, 17, 68, 84). Eden et al. ^85^ measured protein half-lives in the range 0.9–20.5 hr in living human cells, while Dörrbaum et al. ^86^ measured the majority of protein half-lives to lie in the range 1–20 days in rat primary hippocampal, neuron-enriched and glia-enriched cultures. Combining the results of these two studies gives us decay rates in the range 1.44 × 10^−3^– 0.770 hr^−1^. Thus, we assume that the decay rates could lie anywhere in the range 0–0.770 hr^−1^ in the *in vivo* human eye.

**_•_ Permeability of the tear-aqueous interface to R, β_Tear−Aq,r_, and permeability of the aqueous-vitreous interface to R, β_Aq−Vit,r_**

**–** *Ex vivo* porcine eye: these parameters are determined by fitting the porcine version of our mathematical model to our experimental data. We note that these are the only two parameters to be obtained by fitting to experimental data, the permeability of the aqueous-vitreous interface to V/VR/RVR being inferred from its permeability to R and the equivalent human permeability values being inferred from the porcine values (see below), while all other parameters are determined from the literature or directly from our experimental data (or from the model steady-state in the case of *v*_Aqinit_ and *v*_Vitinit_ — see below). In the fitted model, we neglect VEGF (V) and both of its compounds with ranibizumab (VR and RVR), since the measured concentration of ranibizumab in the aqueous and vitreous reaches at least 0.1 pmol ml^−1^ for topical application and almost 5 pmol ml^−1^ for intravitreal administration, while the measured concentration of VEGF never exceeds 3 × 10^−2^ pmol ml^−1^ and is almost always 1 × 10^−2^ pmol ml^−1^ or below (one to two orders of magnitude smaller than the ranibizumab concentrations). Thus, we are only solving (governing) Equations 3, 5, 9 and 13 for the topical treatment case, and Equations 5 and 9 for the intravitreal treatment case. We performed two main fittings. First, for topically applied treatment, we fit for β_Tear−Aq,r_ and β_Aq−Vit,r_ to the mean aqueous ranibizumab data points at 20 min, 40 min and 3.5 hr, neglecting the 1 hr data point since we lack confidence in the measurements at that time point (at which no ranibizumab was detected — see the Experimental results section), and neglecting the vitreal ranibizumab measurements since they take values within the range of the control measurements in the vitreous and are inconsistent with the measured values in the aqueous, given that vitreal concentrations should be lower than those in the aqueous for a concentration gradient to be established, where in fact they are measured to be higher (see the Experimental results section). This gives β_Tear−Aq,r_ = 5.93 × 10^−7^ cm hr^−1^ and β_Aq−Vit,r_ = 0.929 cm hr^−1^. In the second fitting, for intravitreal treatment without CPP, we fit for β_Aq−Vit,r_, taking the mean aqueous and vitreal ranibizumab data points at 20 min as our initial conditions and fitting to the mean aqueous ranibizumab data points at 40 min, 1 hr and 3.5 hr. We take the data points at 20 min as our initial conditions (rather than the calculated vitreal and aqueous concentrations at *t* = 0 hr) since the measured vitreal ranibizumab concentration is smaller than we would expect, based on the dose administered to the vitreous (see the Experimental results section). We do not fit to the ranibizumab data points in the vitreous since these are uninformative, being essentially constant over time. This gives β_Aq−Vit,r_ = 0.577 cm hr^−1^. The topical fit provides a good fit in both cases, and actually does a better job in fitting to the *t* = 3.5 hr data point in the aqueous for intravitreal treatment than the second model fit. Therefore, we consider the topical model fit to be preferable, taking this as our default for the *ex vivo* porcine eye (see the Model fitting section for further details).

**–** *In vivo* human eye: we infer the human values from the porcine values, noting that the permeability of a barrier to a chemical is β = *D*/Δ*x* (cm hr^−1^), ^87^ where *D* (cm^2^ hr^−1^) is the diffusivity of the chemical species within the barrier and Δ*x* (cm) is the width of the barrier. We assume that the diffusivity of ranibizumab within the cornea and suspensory ligaments is the same in porcine and human eyes. The porcine cornea has a mean width (anterior-posterior surface) of about 9.6 × 10^−2^ cm, ^57,62^ while the human corneal width is about 5.2 × 10^−2^ cm. ^53,65^ Scaling accordingly, this gives a default human corneal permeability to ranibizumab of β_Tear−Aq,r_ = (9.6/5.2) × 5.93 × 10^−7^ = 1.10 × 10^−6^ cm hr^−1^. The suspensory ligaments (or zonule) have a width (anterior-posterior) of up to 0.255 cm in humans. ^53^ We were unable to find an equivalent measurement for the porcine eye. Given that porcine and human eye dimensions are similar, we assume the zonule width is the same. Therefore, the human default value for β_Aq−Vit,r_ is assumed to be the same as for the pig, that is 0.929 cm hr^−1^. For the sensitivity analysis, we assume that β_Tear−Aq,r_ and β_Aq−Vit,r_ could vary over a range 10% above and below their default values, giving the ranges 0.990–1.21 × 10^−6^ cm hr^−1^ and 0.836–1.02 cm hr^−1^ respectively.

• **Permeability of the aqueous-vitreous interface to V**/**VR**/**RVR (human and porcine),** β_Aq−Vit,j_ **(for** *j* ∈ {*v*, *u*, *w*}): we infer the permeability of the suspensory ligaments (or zonule) to VEGF and its compounds with ranibizumab (V, VR and RVR) from the fitted permeability for ranibizumab (R, see above); assuming, as for β_Aq−Vit,r_, that the permeabilities do not differ between the human and porcine eye. As noted above, the permeability of a barrier to a given chemical species is directly proportional to the diffusivity of that species (β = *D*/Δ*x*). ^87^ Hutton-Smith et al. ^17^ calculated the diffusivities of V, R, VR and RVR using the Stokes-Einstein relation (at 37°C in physiological saline) and the molecular weights of R and V as found in Ferrara et al. ^66^ and Penn et al. ^67^ respectively (inferring the molecular weights of VR and RVR by adding the weights of R and V). This gave the diffusivities of V, R, VR and RVR as *D*_v_ = 5.11 × 10^−3^ cm^2^ hr^−1^, *D*_r_ = 4.82 × 10^−3^ cm^2^ hr^−1^, *D*_u_ = 3.92 × 10^−3^ cm^2^ hr^−1^ and *D*_w_ = 3.39 × 10^−3^ cm^2^ hr^−1^ respectively. Thus, β_Aq−Vit,v_ = (*D*_v_/*D*_r_)β_Aq−Vit,r_ = 0.985 cm hr^−1^, β_Aq−Vit,u_ = (*D*_u_/*D*_r_)β_Aq−Vit,r_ = 0.755 cm hr^−1^ and β_Aq−Vit,w_ = (*D*_w_/*D*_r_)β_Aq−Vit,r_ = 0.653 cm hr^−1^. For the sensitivity analysis, we assume that β_Aq−Vit,v_, β_Aq−Vit,u_ and β_Aq−Vit,w_ could vary over a range 10% above and below their default values, giving the ranges 0.887–1.08 cm hr^−1^, 0.680–0.831 cm hr^−1^ and 0.588–0.718 cm hr^−1^ respectively.

• **Permeability of the vitreo-retinal interface to R**/**VR**/**RVR,** β_Vit−Ret,j_ (for *j* ∈ {*r*, *u*, *w*}):

**–** *In vivo* human eye: permeabilities of the inner limiting membrane to R, RV and RVR were calculated by Hutton-Smith et al. ^68^ using equations relating a species’ retinal permeability to its hydrodynamic radius derived in Hutton-Smith et al., ^84^ which utilised data from Gadkar et al. ^88^. Values were calculated as β_Vit−Ret,r_ = 6.80 × 10^−4^ cm hr^−1^, β_Vit−Ret,u_ = 6.44 × 10^−4^ cm hr^−1^ and β_Vit−Ret,w_ = 6.23 × 10^−4^ cm hr^−1^. For the sensitivity analysis, we assume that β_Vit−Ret,r_, β_Vit−Ret,u_ and β_Vit−Ret,w_ could vary over a range 10% above and below their default values, giving the ranges 6.12–7.48 × 10^−4^ cm hr^−1^, 5.80–7.08 × 10^−4^ cm hr^−1^ and 5.61–6.85 × 10^−4^ cm hr^−1^ respectively.

**–** *Ex vivo* porcine eye: we set β_Vit−Ret,r_ = 0 cm hr^−1^, β_Vit−Ret,u_ = 0 cm hr^−1^ and β_Vit−Ret,w_ = 0 cm hr^−1^ since there is no choroidal blood flow in the *ex vivo* porcine eye, and hence a concentration gradient will not be maintained to sustain a net flux of ranibizumab and its compounds into the retina. Further, the volume of the retina is over an order of magnitude smaller than that of the vitreous; ^68^ thus, the effect of the flux of ranibizumab and its compounds into the retina is negligible.

**_•_ Rate at which retina contributes V to vitreous, φ_Vit,v_**

**–** *In vivo* human eye: Hutton-Smith et al. ^17^ fitted their 2-compartment (aqueous and vitreous) model to individual patients across Muether and Saunders et al.’s ^75,89,90^ human data. We take their mean fitted value, φ_Vit,v_ = 2.34 × 10^−4^ pmol hr^−1^, as our default, and take the minimum and maximum fitted values, 1.02 × 10^−4^ and 4.46 × 10^−4^ pmol hr^−1^, as our range for sensitivity analysis. We note that in a later paper, Hutton-Smith et al. ^68^ refit this parameter for a 3-compartment model (aqueous, vitreous and retina), such that it contributes VEGF to the retina, rather than directly to the vitreous, obtaining a mean value of 7.29 × 10^−4^ pmol hr^−1^ and a range of 2.50 × 10^−4^ to 1.58 × 10^−3^ pmol hr^−1^. These values are similar to, though larger than the earlier fitted values.

We consider the earlier values to be more relevant for our model, since both our model and that of Hutton-Smith et al. ^17^ neglect the retinal compartment and since the additional VEGF produced in the 3-compartment model is lost to the choroid, rather than contributing to the vitreal supply of VEGF.

**–** *Ex vivo* porcine eye: we set φ_Vit,v_ = 0 pmol hr^−1^ since there is no VEGF production in an *ex vivo*eye.

**_•_ Rate of inflow/outflow to/from tear film, ψ_Tear_**

**–** *In vivo* human eye: the turnover rate of tear fluid in the human eye has been measured to range between 3.0–13.2 × 10^−2^ ml hr^−1^, with a mean value of 7.2 × 10^−2^ ml hr^−1^ ^42^ (see also 37, 40, 43, 45, 78, 80, 91, 92). We do not include evaporative tear loss since the rate of evaporation (approx. 6 × 10^−3^ml hr^−1^ ^92^) is an order of magnitude smaller than the total rate of tear turnover, and since ranibizumab is non-volatile and hence will not be lost through evaporation (with ranibizumab-free fluid lost through evaporation being replaced by ranibizumab-free serous fluid produced by the lacrimal gland).

**–** *Ex vivo* porcine eye: since there is no tear fluid production in the *ex vivo* porcine eye, there is no dilution effect, and hence ψ_Tear_ = 0 ml hr^−1^. As in the human, we neglect evaporative tear loss, noting that visual observation suggests that the majority of the drop remains on the cornea during the course of the experiment.

**_•_ Rate of inflow/outflow to/from aqueous, ψ_Aq_**

**–** *In vivo* human eye: we take ψ_Aq_ = 0.15 ml hr^−1^ with Hutton-Smith et al. ^17,68^ and Missel. ^65^ Hutton-Smith et al. ^17^ take their value from Tasman and Jaeger ^70^ who cite values in the range 0.12–0.15 ml hr^−1^ (across seven studies), while Missel ^65^ takes the mean value from eight studies in normal humans summarised in Table 1 of McLaren, ^69^ which contains values ranging between 6.6 × 10^−2^–0.25 ml hr^−1^. We take this latter interval as our range for sensitivity analysis.

**–** *Ex vivo* porcine eye: since there is no aqueous production (or drainage) in the *ex vivo* porcine eye, there is no dilution effect, and hence ψ_Aq_ = 0 ml hr^−1^.

**_•_ Initial V concentration in aqueous, *v*_Aqinit_**

**–** *In vivo* human eye: we set the initial VEGF concentration equal to its modelled equilibrium value in the absence of ranibizumab, giving a default value of *v*_Aqinit_ = 1.56 × 10^−3^ pmol ml^−1^. Comparing with values in the literature, Saunders et al. ^90^ measured VEGF concentrations in AMD patients to lie in the range 7.5 × 10^−4^–2.8 × 10^−3^ pmol ml^−1^ with a mean value of 1.6 × 10^−3^ pmol ml^−1^, while Wakabayashi et al. ^93^ measured VEGF values in the range 1.13 × 10^−4^–2.82 × 10^−2^ pmol ml^−1^ with a mean value of 2.78×10^−3^ pmol ml^−1^ in patients with proliferative diabetic retinopathy (and without early vitreous haemorrhage), and Hutton-Smith et al. ^17,68^ derive values of ∼ 1.7 × 10^−3^ or ∼ 1.75 × 10^−3^ pmol ml^−1^ as the equilibrium solution to the ranibizumab-free problem. These values are all similar, the measurements from Saunders et al. being the most relevant to the present study since they are from AMD patients. We note that our simulated value is equal to Saunders et al.’s mean value to an accuracy of two significant figures.

**–** *Ex vivo* porcine eye: as noted in the β_Tear−Aq,r_ and β_Aq−Vit,r_ parameter justification section above, we neglect VEGF in porcine simulations.

**_•_ Initial V concentration in vitreous, *v*_Vitinit_**

**–** *In vivo* human eye: we set the initial VEGF concentration equal to its modelled equilibrium value in the absence of ranibizumab, giving a default value of *v*_Vitinit_ = 2.24 × 10^−3^ pmol ml^−1^. Comparing with values in the literature, we note that measured values in humans vary widely, reported mean values ranging between 5 × 10^−6^–9.44 × 10^−1^ pmol ml^−1^ (across healthy, proliferative diabetic retinopathy, subfoveal neovascular membrane, idiopathic full thickness macular hole and AMD patients), ^50,93–98^ with reported values for AMD patients ranging between 5 × 10^−6^–8.48 × 10^−4^ pmol ml^−1^. ^50^ We note that our simulated value is within the range of measured values in humans, and just above the single measured range in AMD patients.

**–** *Ex vivo* porcine eye: as noted in the β_Tear−Aq,r_ and β_Aq−Vit,r_ parameter justification section above, we neglect VEGF in porcine simulations.

• **R dose concentration,** *r*_Dose_: the ranibizumab concentration in both the eye drop and the intravitreal injection is 1 mg ml^−1^ = 10^9^ pg ml^−1^. For our model, we require units of pmol ml^−1^. We can convert to these units by dividing the ranibizumab concentration in pg ml^−1^ by its molecular weight/molar mass ^17,66^ which is 48.35 kDa = 48,350 g mol^−1^ = 48,350 pg pmol^−1^ to give 2.07 × 10^4^ pmol ml^−1^. For the sensitivity analysis, we assume that *r*_Dose_ could vary over a range 10% above and below its default value, giving the range 1.86–2.28 × 10^4^ pmol ml^−1^.

**_•_ Initial R concentration in tear, *r*_Tearinit_**

**–** *In vivo* human eye: when a drop is applied (with volume *V*_Drop_), its contents will rapidly merge with the existing tear film (with volume *V*_TearNorm_) before the excess fluid flows off of the cornea. Assuming that this is the first drop to be applied, the ranibizumab concentration will therefore be 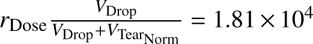, assuming the default value of *V*_Tear_ see discussion of *V*_TearNorm_ above).

**–** *Ex vivo* porcine eye: in this case there is no tear film to dilute the dose since the only fluid on the surface of the cornea is from the drop; therefore, *r*_Tearinit_ = *r*_Dose_ = 2.07 × 10^4^ pmol ml^−1^.

**_•_ Initial R concentration in vitreous, *r*_Vitinit_**

**–** *In vivo* human eye: when an intravitreal injection is administered (with volume *V*_Drop_), it is assumed that the ranibizumab concentration quickly equilibrates throughout the vitreous (with volume *V*_Vit_, given the mixing effect of saccadic eye rotations ^27–31)^, such that 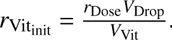 It is assumed that the injected volume does not affect the vitreal volume, the additional volume being negligible (given that *V*_Drop_ « *V*_Vit_) and quickly drained. Therefore, *r*_Vitinit_ = 2.07 × 10^2^ pmol ml^−1^, assuming the default value of *V*_Vit_ (see discussion of *V*_Vit_ above).

**–** *Ex vivo* porcine eye: we use the mean measured value at *t* = 20 min from our experiments for intravitreal ranibizumab injection without CPP, giving a value of *r*_Vitinit_ = 4.29 pmol ml^−1^. Were we to have calculated this parameter in the same way as for the *in vivo* human eye, this would have given a value of *r*_Vitinit_ = 3.00 × 10^2^ pmol ml^−1^ (see the Experimental results and Model fitting sections for a discussion).

## Acknowledgements

This study was funded through the Centre for Systems Modelling and Quantitative Biomedicine by the University of Birmingham Dynamic Investment Fund. P.A.R. was funded by the University of Birmingham Dynamic Investment Fund. C.N.T. was funded by Fight for Sight. G.B.P. was funded by the Kennedy Trust for Rheumatology Research. J.A.R. was funded by the Macular Society. Fig. 1 created in BioRender. Hill, L. (2024) https://BioRender.com/g17g888. Top panel of Fig. 2 created in BioRender. Hill, L. (2024) https://BioRender.com/i25q634.

