## Supplementary Material for "Mathematical Models of Retinal Drug Delivery"

Paul A. Roberts*[1,2], Chloe N. Thomas[3], Gabriel Bellamy Plaice[3], James A. Roberts[3], Marie-Christine Jones[4], James W. Andrews[5] and Lisa J. Hill[3]

[1]Centre for Systems Modelling and Quantitative Biomedicine, University of Birmingham, Birmingham, United Kingdom

[2]Department of Optometry and Visual Sciences, City St George's, University of London, London, United Kingdom

[3]Department of Biomedical Sciences, University of Birmingham, Birmingham, United Kingdom

[4]School of Pharmacy, University of Birmingham, Birmingham, United Kingdom

[5]School of Mathematics, University of Birmingham, Birmingham, United Kingdom

*Corresponding author

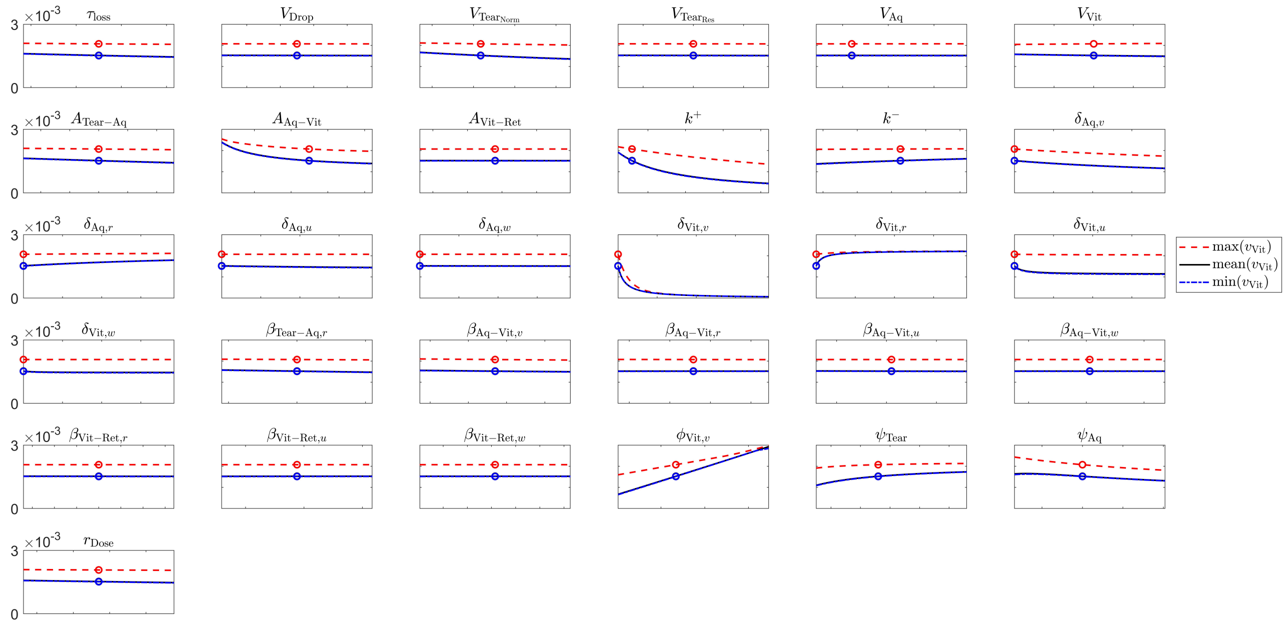


Figure S1: Sensitivity of vitreal VEGF concentration to parameter variation, with topical drop therapy. Panels show variation in the maximum/mean/minimum vitreal VEGF (V) concentration, in response to variation in model parameters over biologically realistic ranges. Equations 2–12 were solved for $t \in [0, 12]$ weeks, with topical drops applied on the hour, every hour. Parameters were varied individually, across 101 values uniformly distributed over the ranges given in Table 5, the remaining parameters being held at their default values given in Table 5. For each parameter set, the maximum/mean/minimum vitreal values of V were calculated over model outputs from the interval $t \in [9, 12]$ weeks. Circles show the maximum/mean/minimum values of V for the default parameter set (see Table 5).

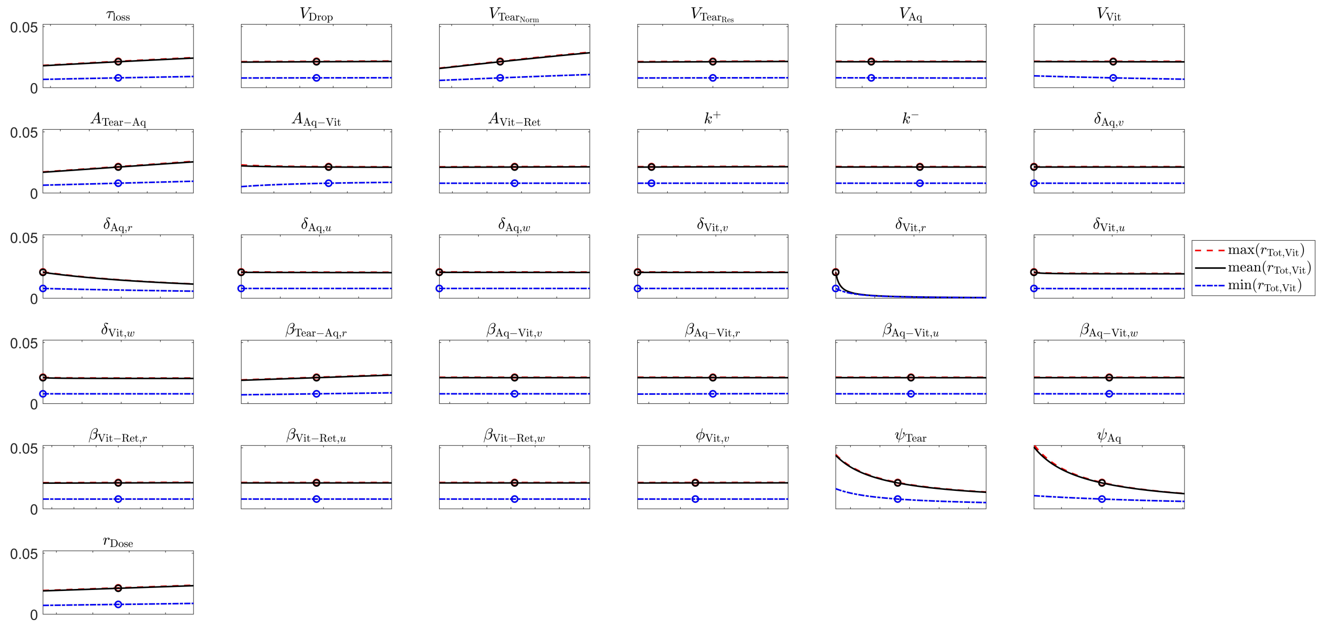


Figure S2: Sensitivity of total vitreal ranibizumab concentration to parameter variation, with topical drop therapy. Panels show variation in the maximum/mean/minimum total vitreal ranibizumab ($R_{Tot}$ = R + VR + 2RVR) concentration, in response to variation in model parameters over biologically realistic ranges. Equations 2–12 were solved for $t \in [0, 12]$ weeks, with topical drops applied on the hour, every hour. Parameters were varied individually, across 101 values uniformly distributed over the ranges given in Table 5, the remaining parameters being held at their default values given in Table 5. For each parameter set, the maximum/mean/minimum vitreal values of $R_{Tot}$ were calculated over model outputs from the interval $t \in [9, 12]$ weeks. Circles show the maximum/mean/minimum values of $R_{Tot}$ for the default parameter set (see Table 5).

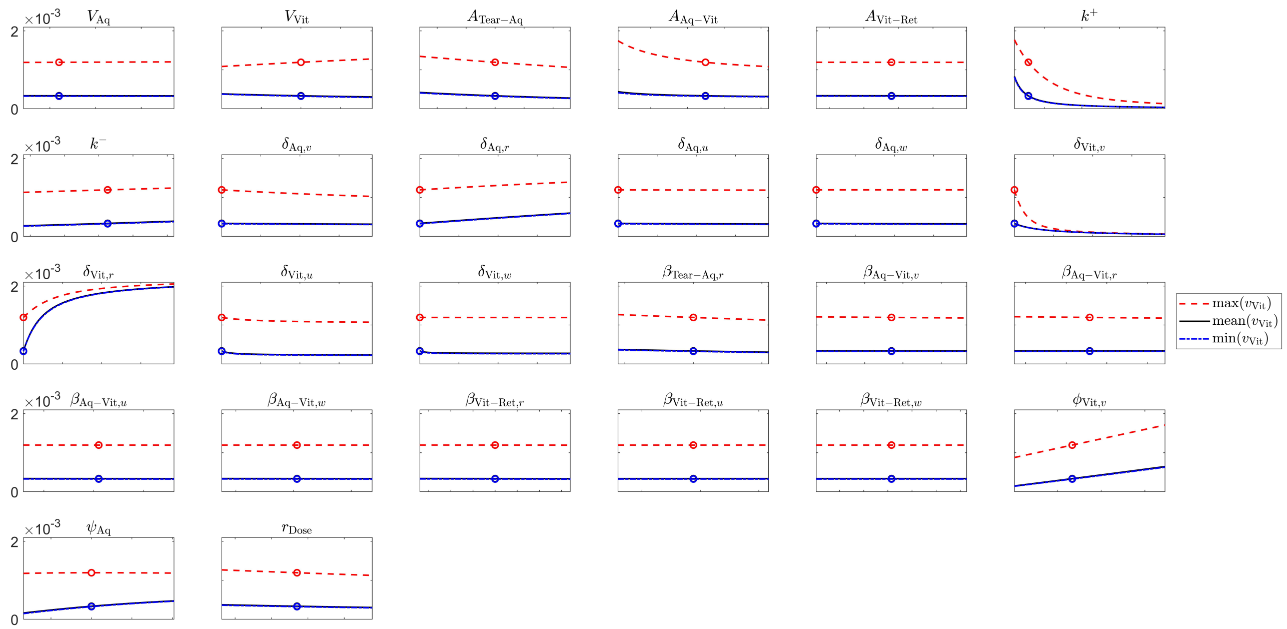


Figure S3: Sensitivity of vitreal VEGF concentration to parameter variation, with drug-eluting contact lens therapy. Panels show variation in the maximum/mean/minimum vitreal VEGF (V) concentration, in response to variation in model parameters over biologically realistic ranges. Equations 2–12 were solved for $t \in [0, 12]$ weeks, with a drug-eluting contact lens, worn continuously. Parameters were varied individually, across 101 values uniformly distributed over the ranges given in Table 5, the remaining parameters being held at their default values given in Table 5. For each parameter set, the maximum/mean/minimum vitreal values of V were calculated over model outputs from the interval $t \in [9, 12]$ weeks. Circles show the maximum/mean/minimum values of V for the default parameter set (see Table 5).

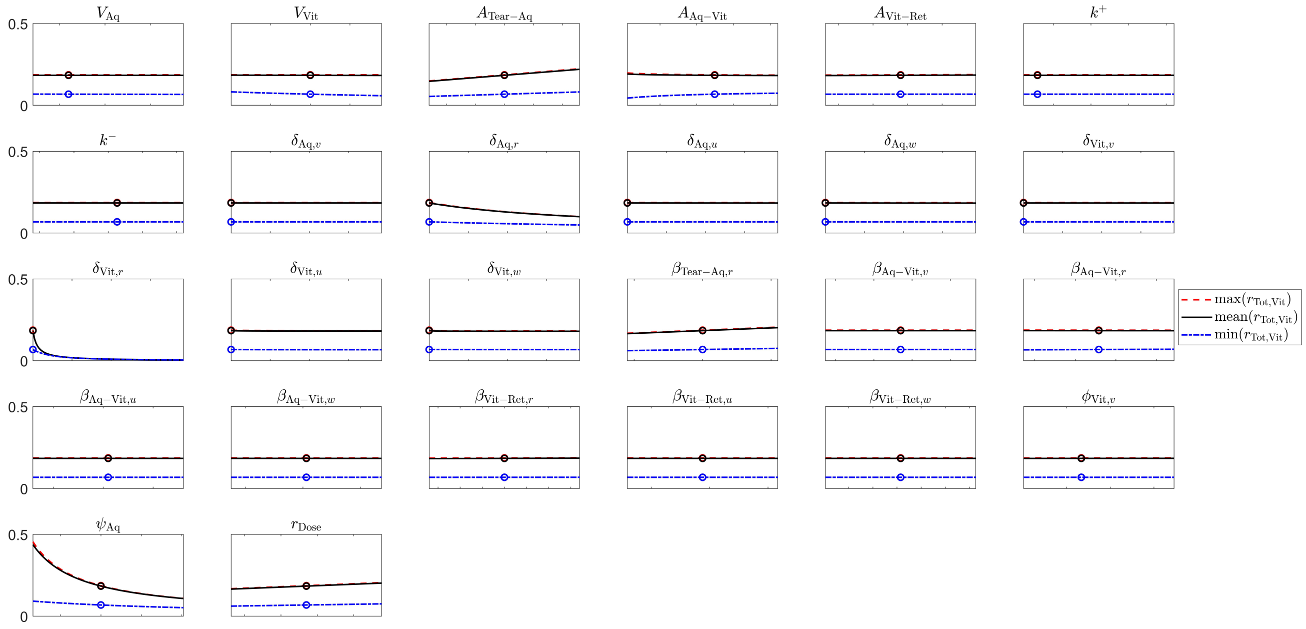


Figure S4: Sensitivity of total vitreal ranibizumab concentration to parameter variation, with drug-eluting contact lens therapy. Panels show variation in the maximum/mean/minimum total vitreal ranibizumab ($R_{Tot}$ = R + VR + 2RVR) concentration, in response to variation in model parameters over biologically realistic ranges. Equations 2–12 were solved for $t \in [0, 12]$ weeks, with a drug-eluting contact lens, worn continuously. Parameters were varied individually, across 101 values uniformly distributed over the ranges given in Table 5, the remaining parameters being held at their default values given in Table 5. For each parameter set, the maximum/mean/minimum vitreal values of $R_{Tot}$ were calculated over model outputs from the interval $t \in [9, 12]$ weeks. Circles show the maximum/mean/minimum values of $R_{Tot}$ for the default parameter set (see Table 5).

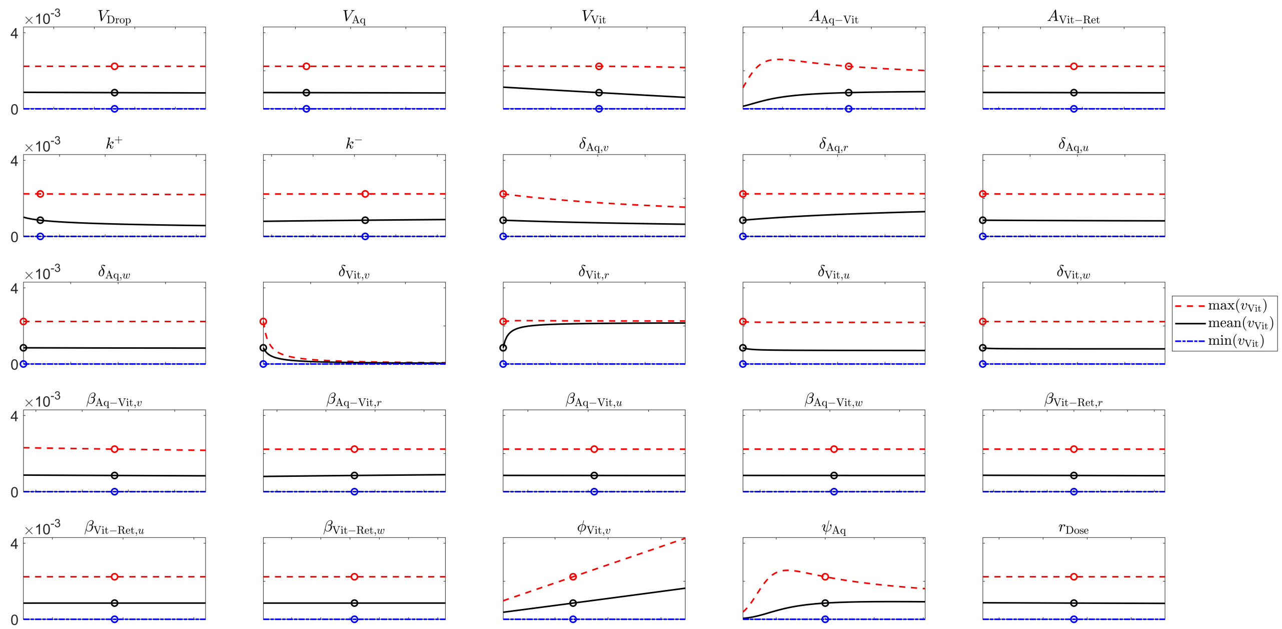


Figure S5: Sensitivity of vitreal VEGF concentration to parameter variation, with intravitreal injection therapy. Panels show variation in the maximum/mean/minimum vitreal VEGF (V) concentration, in response to variation in model parameters over biologically realistic ranges. Equations 2–12 were solved for $t \in [0, 12]$ weeks, with intravitreal injections administered at the start of weeks 1, 5, and 9. Parameters were varied individually, across 101 values uniformly distributed over the ranges given in Table 5, the remaining parameters being held at their default values given in Table 5. For each parameter set, the maximum/mean/minimum vitreal values of V were calculated over model outputs from the interval $t \in [9, 12]$ weeks. Circles show the maximum/mean/minimum values of V for the default parameter set (see Table 5).

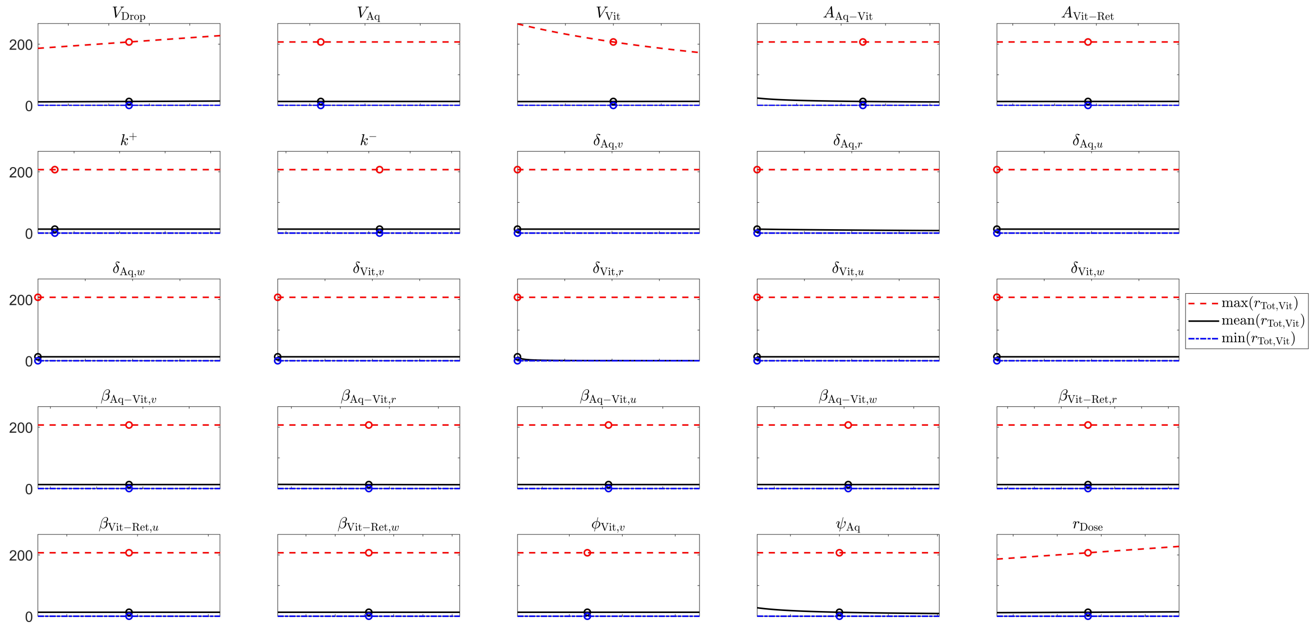


Figure S6: Sensitivity of total vitreal ranibizumab concentration to parameter variation, with intravitreal injection therapy. Panels show variation in the maximum/mean/minimum total vitreal ranibizumab ($R_{Tot}$ = R + VR + 2RVR) concentration, in response to variation in model parameters over biologically realistic ranges. Equations 2–12 were solved for $t \in [0, 12]$ weeks, with intravitreal injections administered at the start of weeks 1, 5, and 9. Parameters were varied individually, across 101 values uniformly distributed over the ranges given in Table 5, the remaining parameters being held at their default values given in Table 5. For each parameter set, the maximum/mean/minimum vitreal values of $R_{Tot}$ were calculated over model outputs from the interval $t \in [9, 12]$ weeks. Circles show the maximum/mean/minimum values of $R_{Tot}$ for the default parameter set (see Table 5).
